# A molecular basis for plant SCAR/WAVE functional divergence

**DOI:** 10.1101/2024.10.01.616141

**Authors:** Sabine Brumm, Aleksandr Gavrin, Matthew Macleod, Guillaume Chesneau, Annika Usländer, Sebastian Schornack

**Affiliations:** Sainsbury Laboratory (SLCU), University of Cambridge, 47 Bateman Street, Cambridge CB2 1LR, UK; Department of Molecular Biology and Genetics, Aarhus University, Aarhus, Denmark; Department of Plant Microbe Interactions, Max Planck Institute for Plant Breeding Research, 50829, Cologne, Germany

## Abstract

Dynamic actin cytoskeleton reorganisation enables plant developmental processes requiring polarised transport such as root hair and leaf trichome formation. The SCAR/WAVE complex plays a crucial role in regulating these dynamics through ARP2/3-mediated actin branching. *SCAR/WAVE* genes occur as small families across a wide range of plant species but whether and how they fulfil different functions remains unclear. We utilise a systematic chimaera approach to define the differential functionality of two closely related *Medicago truncatula* SCAR proteins in plant development. We show that SCAR/WAVE contribution to *Medicago truncatula* root hair or *Arabidopsis thaliana* trichome formation is dependent on two central intrinsically disordered regions (IDRs). Differential functionalities of *Medicago truncatula* SCAR proteins were furthermore associated with the presence/absence of a 42-amino acid sequence within the IDR that impacted protein stability. Through uncovering a molecular basis for functional differences, we advance our understanding of plant SCAR/WAVE complexes.

**One Sentence Summary:** Intrinsically disordered regions in SCAR/WAVE proteins drive diverse functions in root hair and leaf trichome development.

## Introduction

Directional protein transport of endomembrane vesicles along the actin cytoskeleton network is central to plant growth, development and interactions with the environment. In eukaryotes, dynamic actin cytoskeleton organisation is finely tuned by two conserved molecular complexes. The Actin-Related Protein 2/3 (ARP2/3) protein complex controls the formation of branched actin filaments (*1, 2*). Actin branching is pivotal for plant development, and ARP2/3 component mutants display diverse cell polarity related phenotypes (*3 – 6*). ARP2/3 complex activity is regulated by the suppressor of cAMP receptor (SCAR)-WASP family verprolin homologous (WAVE) complex (SCAR/WAVE) consisting of the highly conserved protein components PIR/SRA1, NAP1/NAP125, BRICK1/HSPC300, and one of several ABIL and SCAR proteins. All SCAR proteins have terminal conserved domains, but plant SCAR proteins carry a large and highly variable central region. The N-terminal “SCAR-homology-domain” (SHD) of Arabidopsis thaliana (A. thaliana; At) SCARs physically interacts with ABIL1 and BRK1 (*7, 8*). The C-terminal “Wiskott-Aldrich homology 2, central, and acidic” (WA) domain of plant SCAR proteins activates the ARP2/3 complex (*7 – 9*). Experimental evidence regarding the role and functional conservation of the large variable central region in plant SCAR homologs is still lacking.

Mutations in plant SCAR/WAVE complex components also result in various cell polarity-related phenotypes such as defects in trichome development, cell morphology, division, expansion, and adhesion. The severity of these phenotypes varies depending on whether a single or multi-copy gene complex component is affected. In *A. thaliana*, mutations in the single-copy genes PIR, NAP1, and BRICK1 produce the most severe phenotypes (*10 – 12*), whereas single mutants of the multi-copy ABIL and SCAR subunits exhibit less severe abnormalities (*9, 13, 14*). Of the four *A. thaliana SCAR* (*AtSCAR1-4*) genes, only *scar2* single mutants show the for ARP2/3 pathway related *A. thaliana* mutants characteristic distorted trichome phenotype (*7*). Subsequent genetic studies of *AtSCAR* double, triple and quadruple mutants showed that *AtSCAR1-4* function interchangeably in trichomes but have unequal redundancy (*9, 13*). A threshold model for *A. thaliana* ARP2/3 activation has therefore been proposed in which the individual contribution of the four *At*SCARs varies depending on their biochemical efficiency and expression levels in different cell types, tissues, or organs. However, the precise mechanisms underlying the differences in expression levels and biochemical properties remain unclear.

In *Medicago truncatula* (*M. truncatula; Mt*), *API* (*Aberrant Primordia Invasion*) is the only *SCAR* gene that has been functionally characterised to date (*15*). *MtAPI* is a close homolog of *AtSCAR2. MtAPI* mutants have shorter root hairs, enhanced resistance to Phytophthora infection, and are impaired in root nodule symbiosis with nitrogen fixing bacteria (*16*). Interestingly, the function of *MtAPI* in *M. truncatula* can be substituted by its *A. thaliana* counterpart, *AtSCAR2*. However, it remains uncertain whether *MtAPI* can reciprocally replace the functionality of *AtSCAR2* in *A. thaliana* plants. *M. truncatula* also encodes two additional *Homologs of API* (*HAPI1, HAPI2*) but whether all *M. truncatula SCAR* genes have the same or different functions has not been addressed.

We investigated the two closely related *SCAR* genes *MtAPI* and *MtHAPI1* and their functionality in *M. truncatula* roots and *A. thaliana* leaves. We discovered that *MtHAPI1* cannot functionally replace its close homolog *MtAPI*, as it is unable to complement the *api* mutant. Interestingly, *MtHAPI1* but not *MtAPI* can complement *A. thaliana* defective in *AtSCAR2*. This assigns divergent functions to both *M. truncatula SCAR* genes. Using a chimaera approach, our research revealed the significance of two intrinsically disordered regions within the large central domain of MtAPI and MtHAPI1, emphasising the crucial role of this domain in dictating functionality. Furthermore, a destabilising element present in MtAPI and missing in MtHAPI1 governs SCAR protein abundance in *A. thaliana*. Our findings document how closely related SCAR/WAVE proteins can exert different functionality via variation in their intrinsically disordered domains, advancing our understanding of SCAR/WAVE complex specificity.

## Results

### Paralogous *Medicago truncatula* SCAR/WAVE proteins differ in their functionality

Our previous *M. truncatula* research has shown that *AtSCAR2* and the *Lotus japonicus* homolog *SCARN* can substitute *MtAPI* functionality when expressed in *M. truncatula* roots (*15*) suggesting functional conservation. Phylogenetic analysis shows that *At*SCAR2 is closely related to *Mt*API and *Mt*HAPI1 (Fig. 1A), raising the question whether *Mt*API and *Mt*HAPI1 have similar functions.

**Fig. 1:**
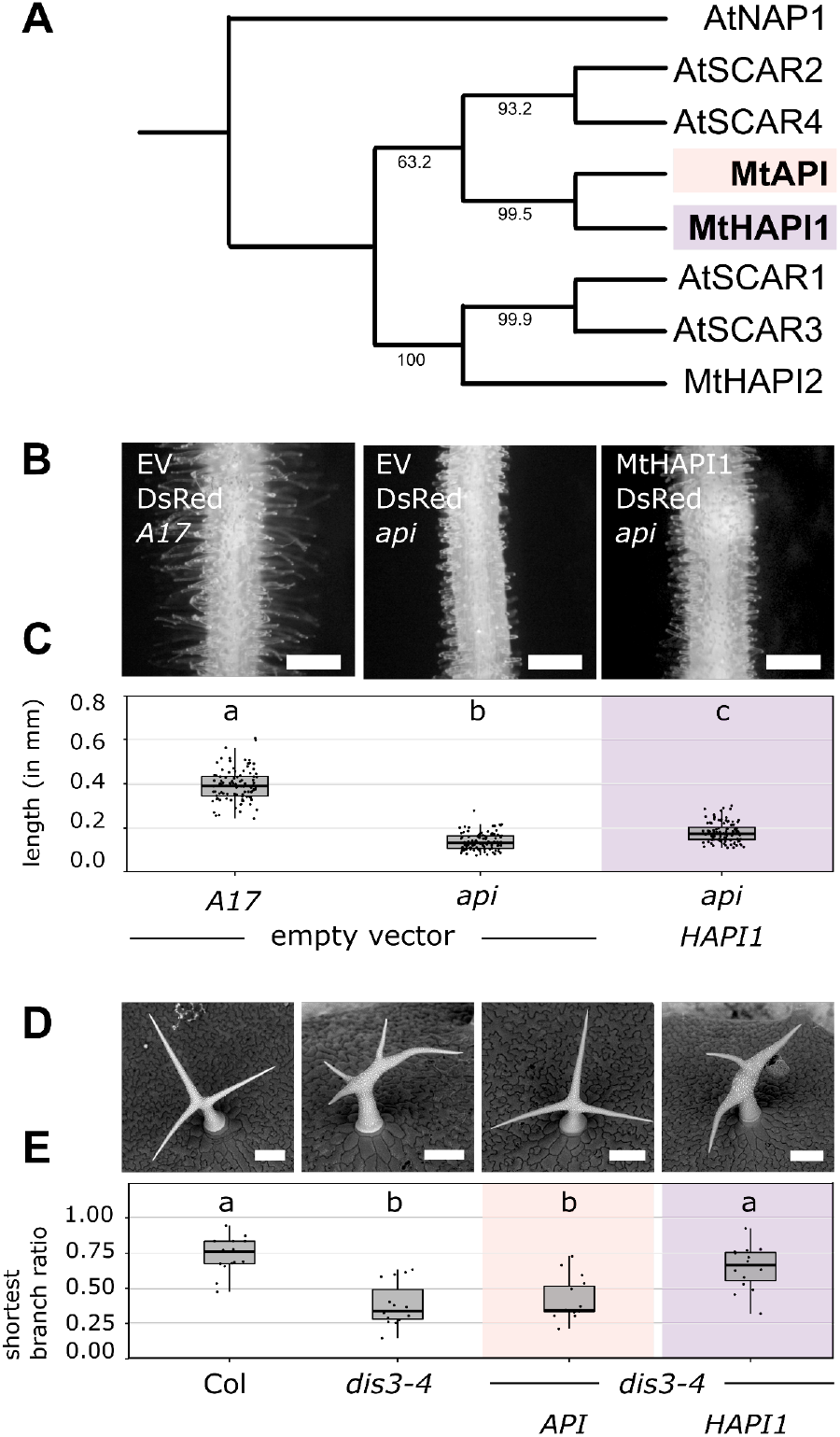
Two closely related *M. truncatula* SCAR proteins exhibit different functions. **(A)** Maximum likelihood phylogenetic tree of *A. thaliana* and *M. truncatula* SCAR proteins with AtNAP1 as outgroup. *M. truncatula* proteins of interest in this study: MtAPI (orange, bold), MtHAPI1 (purple, bold). Node bootstrap values are shown. (**B)** Epifluorescence microscopy images of transgenic *API* and *api M. truncatula* hairy roots expressing an empty *pAtUBQ10:dsRed* vector control or coexpressing *pAtUBQ10:dsRed* with *MtHAPI1* under control of the *MtAPI* promoter. Scale bars, 0.5 mm. (**C)** Root hair length measurements (in millimetres (mm); n = 100 / genotype). Statistics: Shapiro-Wilk test, followed by Kruskal-Wallis with Bonferroni correction; significance differences are indicated by letters a, b and c. (**D)** Scanning electron micrographs of *A. thaliana* trichomes from Col, *dis3-4*, and *dis3-4* lines transformed with *MtHAPI1* or *MtAPI* driven by *AtUBQ3* promoter. Scale bars, 90 µm. (**E)** Shortest trichome branch ratios (n = 15 / genotype). Statistics: Shapiro-Wilk test, followed by Kruskal-Wallis with Bonferroni correction; significance differences are indicated by letters a and b.

To test this we introduced *MtHAPI1* under control of a 2-kb long *MtAPI* promoter sequence (*15*) into roots of *M. truncatula api. MtHAPI1* did not complement the short root hair length phenotype of the *api* mutant (Fig. 1, B and C) while the 2-kb promoter driven *MtAPI* did (*15*). Thus, *MtHAPI1* cannot substitute for the *MtAPI* root hair development function in *M. truncatula*.

Next, we tested whether *MtAPI* and *MtHAPI1* could substitute for the loss of *AtSCAR2/DISTORTED3* in the *A. thaliana dis3-4* mutants (*7*). These plants have normally developed root hairs but display distorted trichome phenotypes. We introduced *MtAPI, MtHAPI1*, and *AtSCAR2*, all under the *A. thaliana UBIQUITIN3* (*AtUBQ3*) promoter, into *dis3-4* (Figure 1, D and E; fig. S1A and B). In this context, *MtHAPI1* and *AtSCAR2* successfully rectified the trichome development defects of *A. thaliana dis3-4* while *MtAPI*-expressing plants developed distorted trichomes (Fig 1, D and E; fig. S1B). Together, these results suggest that *M. truncatula MtAPI* and *MtHAPI1* are phylogenetically close but differ in their functional specificity.

### *Mt*API and *Mt*HAPI1 specificity is encoded within two intrinsically disordered regions

To determine the molecular basis underpinning functional differences of *Mt*API and *Mt*HAPI1, we analysed their protein sequences and systematically assessed the function of API/HAPI1 chimeric proteins (Fig. 2, fig. S2). A comparison of the *Mt*API and *Mt*HAPI1 protein sequences revealed five distinct regions of varying sequence similarity. The first, third, and fifth regions exhibited high sequence conservation, while the second and fourth regions displayed greater variability including amino acid segments missing in *Mt*HAPI1 (fig. S2A). The IUPred3a web server (https://iupred3.elte.hu/) predicts a high probability for intrinsic disorder (scores between 0.5 - 1) in regions 1 (post-SHD domain), 2, 4, and 5 (fig. S2B and C). Given the conservation of the SHD and WA domains in regions 1 and 5, associated with SCAR/WAVE complex interaction and ARP2/3 activation (*7 - 9*), respectively, we hypothesised that variation within regions 2 and 4 may account for the differential specificity.

**Fig. 2:**
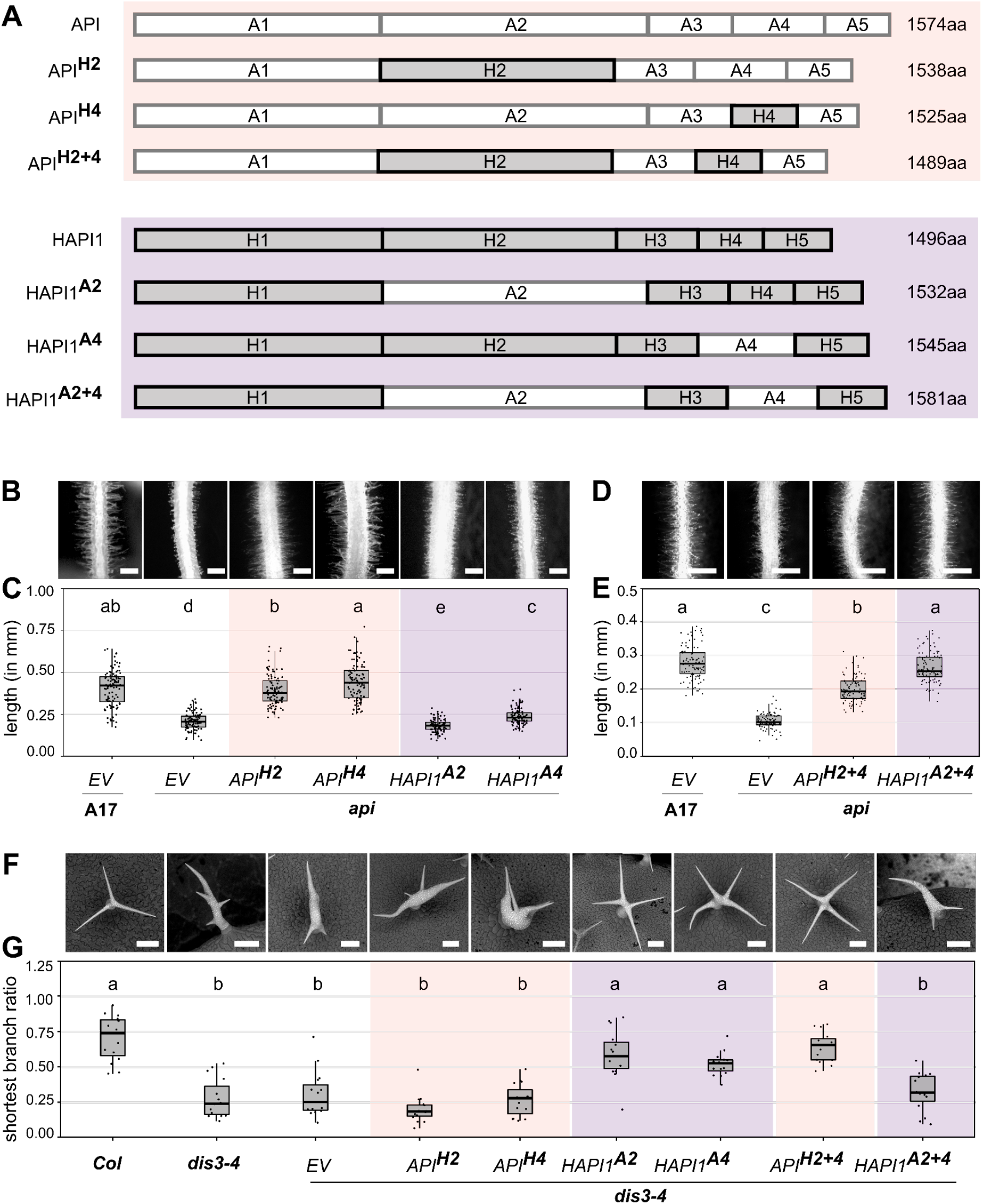
Exchanging two regions between *Mt*API and *Mt*HAPI1 switches their functions. **(A)** Schematic of *Mt*API (orange) and *Mt*HAPI1 (purple) chimeric proteins, with *Mt*API regions in white and *Mt*HAPI1 in grey. (**B** to **E)** Epifluorescence microscopy images of transgenic *M. truncatula API (A17)* and *api* hairy roots expressing an empty *pAtUBQ10:dsRed* vector control or coexpressing *pAtUBQ10:dsRed* with chimeric *MtAPI/HAPI1* variants (region 2 or 4 exchanged in (B); both in (D)) under the *MtAPI* promoter. Scale bars, 0.5 mm. Root hair length measurements of *api* roots with single region (C) or double region (E) *MtAPI/MtHAPI1* chimaeras in millimetres (n = 100 / genotype). Statistics: Shapiro-Wilk test, followed by Kruskal-Wallis with Bonferroni correction; significance differences are indicated by letters a, ab, b, c, d, and e **F)** Scanning electron micrographs of *A. thaliana* trichomes from untransformed Col, *dis3-4* and *dis3-4* lines expressing *empty vector* (EV) or chimeric *MtAPI/HAPI1* variants under the *AtUBQ3* promoter. Scale bars, 90 µm. **G)** Shortest trichome branch ratios (n = 15 / genotype). Statistics: Shapiro-Wilk test, followed by Kruskal-Wallis with Bonferroni correction; significance differences are indicated by letters a and b. Orange: *MtAPI* chimaeras; Purple: *MtHAPI1* chimaeras.

To identify regions with functional specificity in *M. truncatula* and *A. thaliana*, we exchanged region 2, region 4, or both regions between the two paralogs (Fig. 2A). Expression of the resulting chimeric proteins under the *MtAPI* promoter in *M. truncatula api* roots revealed that swapping a single region between *Mt*API and *Mt*HAPI1 was insufficient to alter their functionality (Fig. 2, B and C; fig. S2D). However, when regions 2 and 4 were swapped together, *Mt*API failed to confer normal root hair development having significantly shorter root hairs, while *Mt*HAPI1 gained functionality in root hair development (Fig. 2, D and E; fig. S2D). Hence, *Mt*API regions 2 and 4 both contribute to its functionality in *M. truncatula* root hair development.

We next analysed the chimaeras for their ability to rescue the trichome defects of the *A. thaliana dis3-4* mutant. Exchanging a single region between *Mt*API and *Mt*HAPI1 did not alter their functionality (Fig. 2, F and G; fig. S2D). However, the simultaneous exchange of regions 2 and 4 compromised *Mt*HAPI1 functionality in *A. thaliana*, whereas comparable *Mt*API chimaeras rescued the trichome development defects of the *dis3-4* mutant defective in *AtSCAR2*. We therefore conclude that *Mt*HAPI1 regions 2 and 4 together condition function in *A. thaliana* trichome development.

Taking the data from *M. truncatula* and A. thaliana together, we demonstrated that exchanging regions 2 and 4 between MtAPI and MtHAPI1 exchanges their functionality.

### Functional specificity of *Mt*SCARs in *Arabidopsis thaliana* is determined by specific amino acid residues rather than sequence length

To better understand how SCAR sequence variation contributes to the distinct molecular functionalities of *Mt*API and *Mt*HAPI1, we generated an extended phylogenetic tree of SCAR proteins derived from 9 different legume genomes (fig. S3A). To further curate our comparison, we focused solely on *Mt*API and *Mt*HAPI-like protein sequences and removed all *Mt*HAPI2-like sequences from subsequent alignments. This comparative analysis revealed that all *Mt*HAPI1-like proteins lack a 42-amino-acid segment at the intersection of *Mt*API region 1 and 2 (Segment A), as well as two segments of 20 (Segment B) and 29 amino acids (Segment C) in *Mt*API region 4 (fig. S3B), resulting in an overall shorter length. Some amino acid residues within these API-like segments are highly conserved, others vary significantly between the different species (fig. S4).

To investigate whether differential *Mt*API and *Mt*HAPI1 functionality is influenced by their overall length or protein sequence composition, we either swapped all identified amino acid segments between *Mt*API and *Mt*HAPI1, or substituted them with the Glycine/Serine linkers of the same length (Fig. 3A). An mCitrine tag introduced into a permissive site (*17*) within region 2 of all *Mt*API and *Mt*HAPI1 variants allowed us to also assess localisation and abundance of these protein variants.

**Fig. 3:**
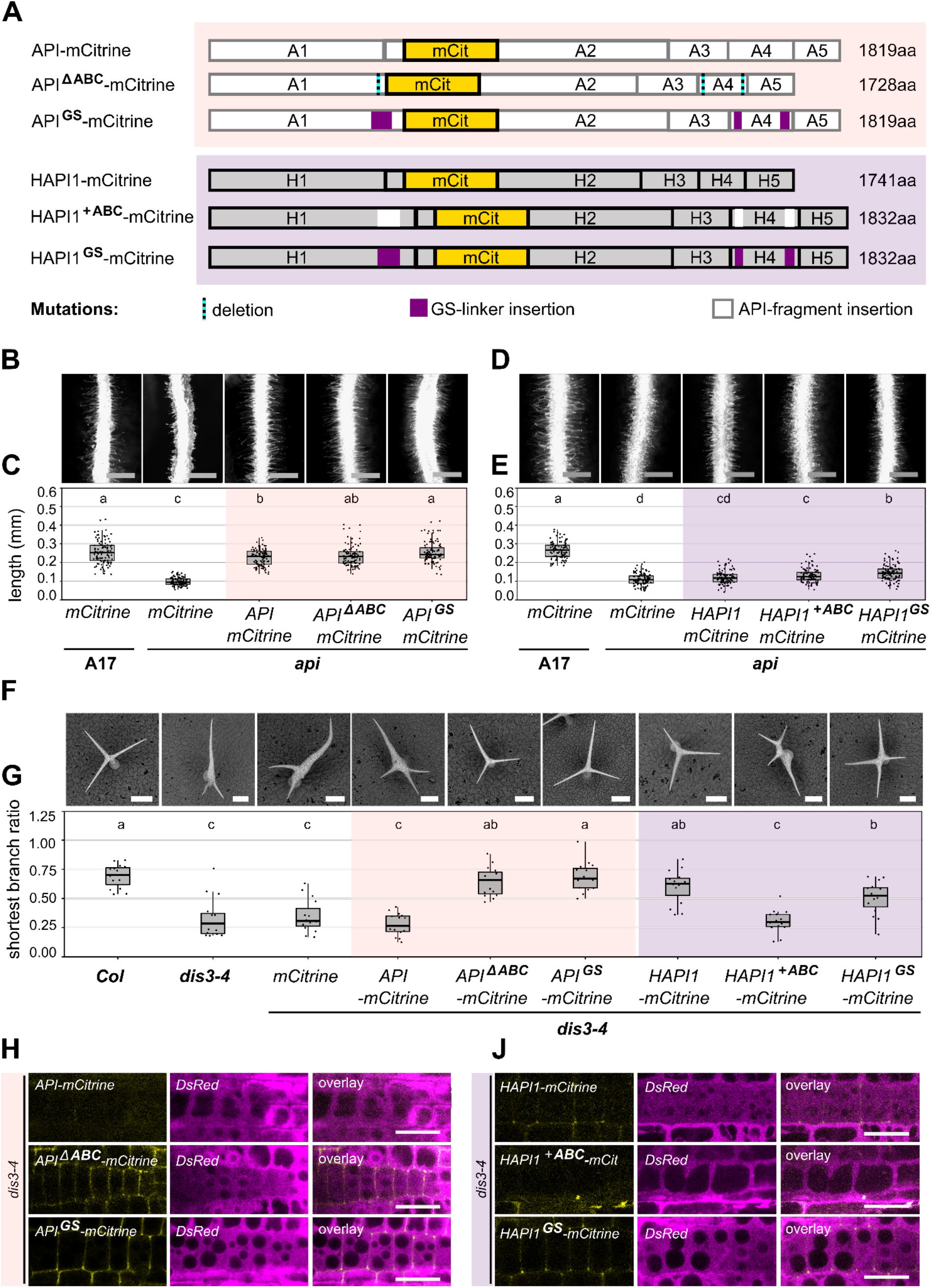
Three *Mt*API amino acid segments contribute to functional separation. **(A)** Schematic of *Mt*API (orange) and *Mt*HAPI1 (purple) segment A, B and C mutant protein variants with fluorescent tags. Segments A, B and C in *Mt*API are either deleted (dotted lines on cyan) or replaced with GS-linkers (magenta). In *Mt*HAPI1, *Mt*API segments A, B and C (white) or GS-linkers are inserted. mCitrine tags are shown in yellow, with protein size changes indicated. (**B** to **E)** Epifluorescence microscopy images of *M. truncatula A17* and *api* roots, coexpressing *MtAPI* (B) *or MtHAPI1* (D) segment mutants with *pAtUBQ10:dsRed*. Scale bars, 0.5 mm. Root hair length measurements of MtAPI (C, orange) or MtHAPI1 (E, purple) segment mutants in *api* (in millimetres (mm), n = 100 / genotype). Statistics: Shapiro-Wilk test, followed by Kruskal-Wallis with Bonferroni correction; significance differences are indicated by letters a, b, c and d. (**F)** Scanning electron micrographs of *A. thaliana* trichomes from *Col, dis3-4*, and *dis3-4* lines expressing *MtAPI* or *MtHAPI1* segment variants under the *AtUBQ3* promoter. Scale bars, 90 µm. (**G)** Shortest branch ratios (n = 15 / genotype). Statistics: Shapiro-Wilk test, followed by Kruskal-Wallis with Bonferroni correction; significance differences are indicated by letters a, b and c. (**H** and **J)** Confocal microscopy of *Mt*API (H) and *Mt*HAPI1 (J) segment mutant expression and localisation (yellow) in *A. thaliana* root cells. *DsRed* (magenta) serves as transformation control and cytosolic marker. Scale bars, 20 *μ*m.

First, we tested whether *Mt*API protein segments A, B, and C are essential for *Mt*API functionality in *M. truncatula* root hairs. To that end, we expressed the deletion and GS-linker constructs under control of the *MtAPI* promoter in roots of *M. truncatula api* composite plants (Fig. 3, B to E; fig. S5). As expected, a wildtype API-mCitrine fusion construct complemented the short root hair phenotype of *api* (Fig. 3, B and C). Removing MtAPI segments A, B and C (API^ΔABC^-mCitrine) or replacing these segments with Glycine/Serine (GS)-linkers in *Mt*API (API^GS^-mCitrine) did not impair *Mt*API functionality (Fig. 3, B and C). Inserting *Mt*API segments A, B and C or GS-linkers into *Mt*HAPI1 (HAPI^+ABC^-mCitrine; HAPI^GS^-mCitrine) did not confer the ability to rescue *api* root hair defects (Fig. 3, D and E). We therefore conclude that *Mt*API segments A, B and C do not define *Mt*API functionality and that other regions of the large central domain likely condition functionality in *M. truncatula* roots.

We next tested the same constructs in the context of *A. thaliana* trichome development (Fig. 3, F and G; fig. S5). While *Mt*API does not normally complement distorted trichomes in *A. thaliana dis3-4* mutants, the replacement of *Mt*API segment A, B, and C with GS-residue linkers (API^GS^-mCitrine) or their complete deletion (API^ΔABC^-mCitrine) resulted in a gain of function, similar to *Mt*HAPI or *At*SCAR2 (Fig. 3, F and G). API^delABC^-mCitrine and API^GS^-mCitrine polarly localised to the plasma membrane in *A. thaliana* root cells, with visible signal accumulation at the cell periphery (Fig. 3H), similar to the known localisation of *At*SCAR1 (*18*). By contrast, no API-mCitrine signals were visible at the cell periphery. Unlike wild-type *Mt*HAPI, the insertion of *Mt*API amino acid segments A, B, and C into *Mt*HAPI1 (HAPI^+ABC^-mCitrine) failed to rescue the *A. thaliana dis3-4* trichome phenotype (Fig. 3, F and G). Interestingly, the substitution of these amino acids with the same number of GS-residues (HAPI^GS^-mCitrine) maintained functionality. In contrast to API-mCitrine, weak HAPI1-mCitrine signals were detectable at the cell peripheries (Fig. 3J). The insertion of GS-linkers did not interfere with HAPI1-mCitrine fluorescence, but when we inserted *Mt*API segments A, B and C into HAPI1-mCitrine, we were no longer able to detect fluorescence signals. Thus, the presence of *Mt*API segments A, B and C abolishes trichome functionality and diminishes peripheral fluorescence signals. This suggests that three specific amino acid segments in the *Mt*API regions 2 and 4 control specificity, protein levels and localisation in *A. thaliana*.

### A 42-amino acid sequence of *Mt*API destabilises proteins in *Arabidopsis thaliana* and

#### Nicotiana benthamiana

To determine the amino acid segments in *Mt*API region 2 and 4 which prevent complementation of *A. thaliana dis3-4*, we tested additional *Mt*API deletion constructs (Fig. 4, A to C; fig. S6A). API^ΔA^-mCitrine significantly restored trichome morphology, albeit not to full extent, while API^ΔB^-mCitrine and API^ΔC^-mCitrine were indistinguishable from the mCitrine negative control. API^ΔAB^-mCitrine and API^ΔAC^-mCitrine fully rescued trichome morphology, while API^ΔBC^-mCitrine significantly but not fully increased the length of the shortest trichome branch compared to the negative controls. Taking the data from fluorescence imaging and trichome morphology together, we conclude that the 42-amino acid sequence of segment A is sufficient to prevent restoration of trichome morphology and to lower *Mt*API protein abundance.

**Fig. 4:**
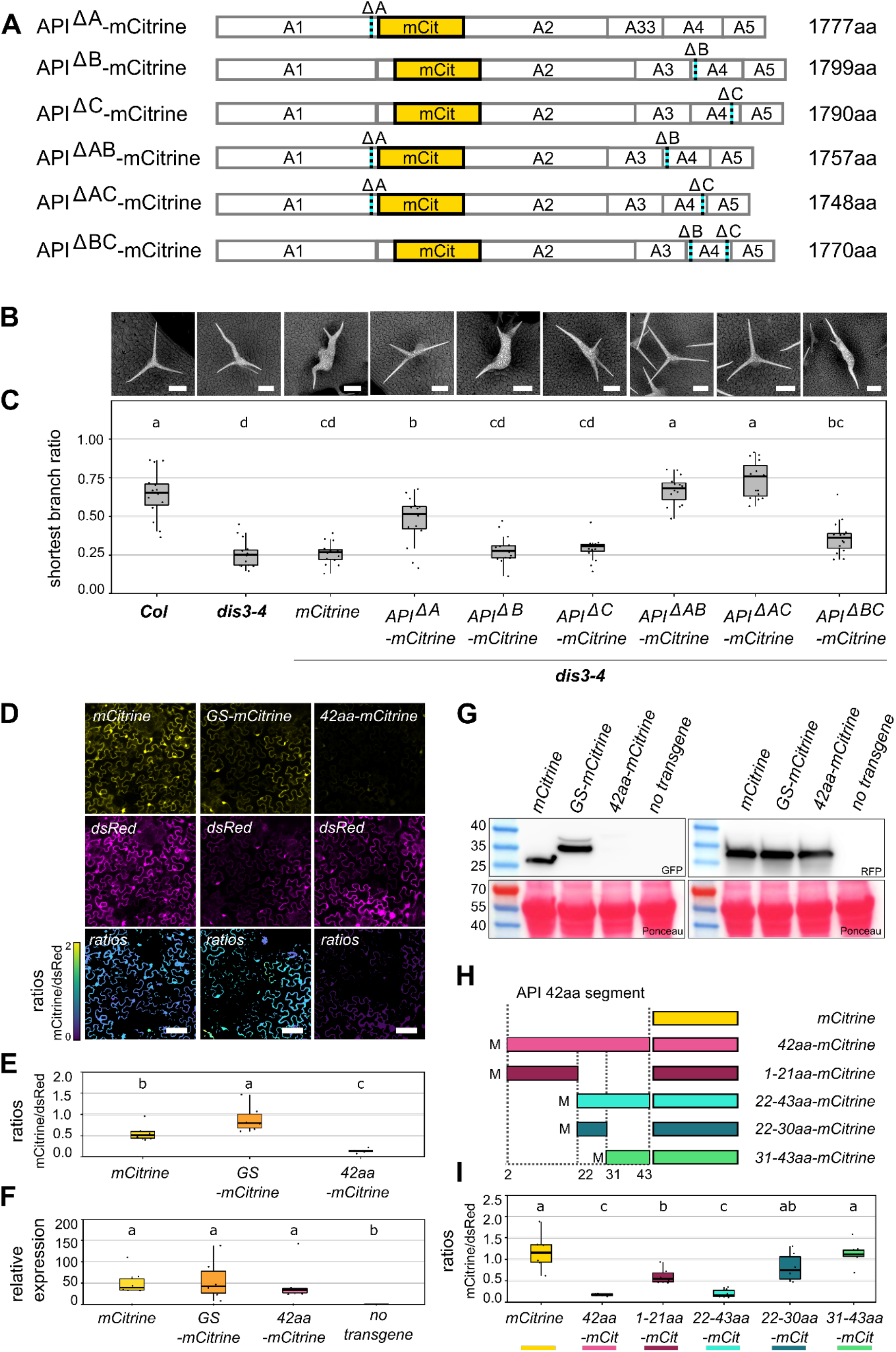
*Mt*API segment A acts as a destabilising element in *A. thaliana* and *N. benthamiana*. (**A**) Schematic of *Mt*API protein variants with single or double deletions in segments A, B, and C (dotted lines over cyan). mCitrine tags are shown in yellow, with protein size changes depicted. (**B**) Scanning electron micrographs of *A. thaliana* Col, *dis3-4*, and *dis3-4* lines expressing *Mt*API segment A, B and C deletion variants under the control of the *AtUBQ3* promoter. Scale bars, 90 µm. (**C**) Shortest trichome branch ratio (n=15/genotype). Statistics: Shapiro-Wilk test, followed by Kruskal-Wallis with Bonferroni correction; significance groups: a, b, bc, cd, d (**D**) Confocal images of mCitrine, GS-mCitrine, and 42aa-mCitrine in *N. benthamiana* pavement cells (yellow) with dsRed control (magenta). Z-projections of fluorescence ratios shown as false color images. Scale bars, 100 *μ*m. (**E**) Fluorescence quantification relative to dsRed. Statistics: Shapiro-Wilk test, followed by Kruskal-Wallis with Bonferroni correction; significance difference groups indicated by letters a, b, c (**F**) Quantification of mCitrine variants mRNA levels in *N. benthamiana* leaves, using 2-ΔΔCP with *NtL23* and *NtFBox* as reference genes (n=6). Negative control: untransformed leaf samples. Statistics: Shapiro-Wilk test, followed by Kruskal-Wallis with Bonferroni correction; significance groups: a, b. (**G**) Western Blot membrane cut-outs (see Suppl. Fig. 6b,c for whole membranes) of mCitrine, GS-mCitrine, 42aa-mCitrine, API-mCitrine and coexpressed dsRed protein levels in transiently transformed *N. benthamiana* leaves. Membranes were stained with Ponceau and probed with GFP or RFP antibodies. Expected protein sizes: ponceau stained Rubisco (55kDa), dsRed (28kDa); mCitrine (27kDa); GS-mCitrine (36kDa), 42aa-mCitrine (38kDa). Prestained protein ladder: Page Ruler 10-180kDa. (**H**) Colour coded schematic of 42aa-mCitrine deletion constructs. Deletion parts are confined by dotted lines, followed by mCitrine tag. (**I**) Fluorescence quantification: mCitrine /dsRed ratios were calculated using the FIJI plugin FRETENATOR (n=6). Statistics: Shapiro-Wilk test, followed by Kruskal-Wallis with Bonferroni correction; significance difference groups indicated by letters a, b, c. Construct colour code refers to schematic (H).

We next tested whether the 42-amino acid sequence of *Mt*API segment A functions independent of the SCAR protein or plant species context. In *Nicotiana benthamiana* (*N. benthamiana*) transient expression experiments under *A. thaliana UBIQUITIN3* promoter control, the fluorescent signal was significantly reduced in 42aa-mCitrine fusions compared to mCitrine alone (Fig. 4, D and E). Replacing the 42-amino acids with a GS-linker of equal length did not reduce mCitrine fluorescence levels. While the transcript levels of mCitrine and 42aa-mCitrine were similar, protein levels differed significantly (Fig. 4, F and G; fig. S6, B and C). Thus, the addition of a 42 amino acid sequence contained within *Mt*API segment A reduces protein abundance across different proteins, species and tissue contexts.

Protein stability can be affected by post translational modifications, including glycosylation, ubiquitination and phosphorylation. The 42-amino acid sequence contains several potential post translational modification sites (fig. S7B). A glycosylation site was predicted with high confidence for asparagine at position 5 (513 in *Mt*API). Deletion of the asparagine residue in the 42-amino acid mCitrine construct (41aa-mCitrine) reduced mCitrine fluorescence even further (fig. S7, B and C). Mass spectrometric analysis of immunoprecipitated 42aa-mCitrine revealed a ubiquitinated lysine residue at position 12 (K520 in *Mt*API) (fig. S7B). However, substituting all lysine residue (9, 10, 12 and 39) to arginine (R) residues (4xKtoR-mCitrine) did not restore mCitrine fluorescence to control levels (fig. S7, B and D). Substituting the corresponding K residues to R in API-mCitrine (K517R, K518R, K520R and K547R) did not lead to a gain of function of *Mt*API in *A. thaliana* (fig. S8, A to C). Neither the single, nor multiple substitution mutant API-mCitrine variants were able to rescue the *A. thaliana dis3-4* distorted trichome phenotype suggesting that lysine-targeted ubiquitination of the 42-amino acid sequence does not contribute to *Mt*API destabilisation in *A. thaliana*.

Further dissection of the peptide into four parts (1-21aa, 22-43aa, 22-30aa and 31-43aa) (Fig. 4H; fig. S7B) and their expression as mCitrine fusions in *N. benthamiana* leaves showed that amino acids 22-43 drove the most significant reduction of mCitrine fluorescence levels (Fig. 4I). These 22 amino acids include four phosphorylated serine and threonine residues (S22, T23, T28, S30) (fig. S7B). To abolish phosphorylation or mimic constitutive phosphorylation, we changed them to alanine (A) or aspartic acid (D) residues, respectively, in 42aa-mCitrine (4xSTtoA- and 4xSTtoD-mCitrine) (fig. S7, B and E). None of these modifications resulted in fluorescence levels comparable to mCitrine alone (fig. S7E). We conclude that phosphorylation of the 42-amino acids does not contribute to the decreased fluorescence levels of 42aa-mCitrine. In summary, the results demonstrate that a segment of 42-amino acids lowers protein abundance of *Mt*API and other proteins in *A. thaliana* and *N. benthamiana* via an unidentified mechanism.

## Discussion

In this study, we show that two closely related *M. truncatula* SCAR homologs underwent functional diversification following a duplication event in leguminous plants. SCAR/WAVE proteins are part of the evolutionary conserved actin regulatory SCAR/WAVE complex in eukaryotes. While some components of this complex have been retained as single copy genes during evolution, other components, such as the *SCAR* genes, have undergone duplication and sequence diversification across multiple species. The diversification raises intriguing questions about potential functional differences of SCAR/WAVE subcomplexes. Interestingly, recent results of investigations of the SCAR/WAVE downstream target, the ARP2/3 complex, have suggested the existence of functionally diverse ARP2/3 subcomplexes with varying subunit compositions (*19*). It is thus possible that different ARP2/3 subcomplexes, with varying subunit compositions, could selectively interact with distinct SCAR/WAVE subcomplexes. Such interactions may allow for greater diversity in actin regulation across various cellular contexts.

Although previous studies have identified SCAR mutants with visible phenotypes in *Oryza sativa* (*20*), *Lotus japonicus* (*21*) and *M. truncatula* (*22*), systematic functional studies on SCAR diversification have only been performed in *A. thaliana*. Our study expands this knowledge by showing that two *M. truncatula* SCAR proteins are not functionally interchangeable in *M. truncatula* or *A. thaliana* (Fig. 1). This contrasts with *A. thaliana* SCARs which function interchangeably in trichome development when expressed under the same regulatory elements (*13*). To rule out a contribution of differential promoter activities among the different variants, all constructs in our study were driven under the same promoter. Furthermore, the chimeric proteins with domain swaps did retain their original activation domains (WA).

This study explores the functional similarities and differences of *MtAPI* and *MtHAPI1* in *M. truncatula* roots and *A. thaliana* leaves. We have not directly compared *MtHAPI1* with other previously assigned functions of *MtAPI* in *M. truncatula*, such as its role in susceptibility to oomycete root infection (*15*) or its requirement for nitrogen-fixing symbiosis (*16*). Future studies could benefit from the use of new genome editing tools to further explore *MtHAPI1* and *MtHAPI2* roles more comprehensively since there are no *M. truncatula* mutants available yet. However, our findings suggest a potential role for *MtHAPI1* in trichome development, a function distinct from that of *MtAPI*. While *MtAPI* mutants do not exhibit trichome defects (*15*), the *rit1* mutant, which harbours a defective allele of *M. truncatula NAP1* - a single-copy SCAR/WAVE complex member - displays both root hair and trichome defects (*11*). Since *MtAPI* is not involved in trichome development, it is likely that this function may be mediated by *MtHAPI1, MtHAPI2*, or both, although further work is needed to confirm this hypothesis. These findings align with the idea that different SCAR/WAVE subcomplexes could serve tissue-specific roles, with *MtHAPI1* potentially involved in processes related to trichome formation.

One of our key findings shows that functional differences between MtAPI and MtHAPI1 are determined by two IDRs within the large central variable domain, rather than the SHD or WA domains (Fig. 2; fig. S2). This highly variable central region, distinct in plant SCARs (22), represents an area for further investigation, as it may contribute to the unique functionalities observed in legume species. IDRs can impact protein stability through both ubiquitin-dependent and -independent degradation by the proteasome (*23, 24*). The overall structure, rather than a specific primary sequence, often defines this degradation function. It is therefore not surprising that the amino acid sequences contributing to SCAR specificity can be variable. Rather than their sequence, it is the striking SCAR clade specific absence or presence of the destabilising segment in the IDR of several legume species (Fig. S4) that suggests a functional diversification of SCAR homologs across species of the legume clade.

Interestingly, we have identified a 42-amino acid segment within the central region of plant SCAR proteins as a major driver of the differential specificity of MtAPI/HAPI1 (Fig. 3 and 4). Even a shorter 22-amino acid subsegment functions as a destabilising element, impacting SCAR protein abundance (Fig. 4I). Our findings open new avenues for future research to investigate how plants regulate SCAR protein abundance, which does not appear to be controlled by transcript levels or ubiquitination in this case (Fig.4, Fig. S7 and S8). Our results align with previous findings in *A. thaliana*, where SCAR/WAVE complex components transiently accumulate at the root hair initiation site, with abundance inversely correlated to root hair elongation speed (*17*). Although the specific segment we identified in *MtAPI* is not conserved in *AtSCAR2*, similar destabilising elements may exist, providing a potential mechanism for rapid protein dissipation during root hair elongation.

In conclusion, this work has uncovered IDRs as the molecular basis for functional differences between two *M. truncatula* SCAR proteins, advancing our understanding of SCAR/WAVE complex specificity in plant development and plant-microbe interactions. These findings generate exciting new directions for future research, particularly in identifying the biochemical mechanisms of SCAR protein abundance and functional regulation.

## Material and Methods

### Plant materials and Growth Conditions

#### Arabidopsis thaliana

Seeds of *A. thaliana Col-0* and *dis3-4* seeds were kindly provided by Daniel Szymanski. *A. thaliana* plants used for transformation, complementation assays and propagation were grown in Levington F2 soil. The plants were maintained under constant light conditions (170µmol m^−2^s^−1^ PAR) at 21°C (day/night) temperature and 65% relative humidity. For confocal microscopy experiments, seedlings were sterilised with chlorine gas over night and grown on ½ MS pH 5.7 medium (0.22% [wt/v] MURASHIGE & SKOOG MEDIUM (Duchefa Biochemie), 0.05% [wt/v] MES monohydrate (MELFORD) and 1.2% [wt/v] Plant Agar (Duchefa Biochemie).

#### Medicago truncatula

Seeds of *M. truncatula* Jemalong A17 and *api* were propagated from previously described materials (*15*). For general propagation, *M. truncatula* plants were grown under long-day conditions (350µmol m^−2^s^−1^ PAR, 21°C for 16 hrs; 0µmol m^−2^s^−1^ PAR, 17°C for 8 hrs) at 65% humidity. The growth substrate used consisted of 45% Levington F2 soil, 45% Terra-Green stones/sand mixture (50:50) and 10% Perlite. For hairy root transformations, *M. truncatula* plants were grown under long-day conditions with 16 hours of light at 20°C followed by 8 hours of darkness at ambient humidity.

#### Nicotiana benthamiana

For transient leaf infiltration assays and propagation, seeds of *N. benthamiana* were grown on Levington F2 soil under greenhouse conditions. The seeds are progenies of a laboratory cultivar from The Sainsbury Lab, Norwich, UK and originated from Australia (*25*).

### Design of constructs and cloning

Primer design, sequence assembly, and analysis were performed using CLC Main workbench 20. Coding sequences for untagged *MtAPI* (*Medtr4g013235/MtrunA17_Chr4g0004861*) and *MtHAPI1* (*Medtr7g071440/MtrunA17_Chr7g0244031*) were synthesised in *pUC57* by GENEWIZ, Inc. Chimeric *MtAPI* and *MtHAPI* coding sequences (CDS) were generated via Gibson assembly (see Fig. 2A). *MtAPI* and *MtHAPI1* regions were amplified using designated primers with overhangs for assembly: *<u>MtAPI</u>* <u>region 1</u> (*MtAPI* bases 1 to 1548), primer pl_apiSHD_F_AG and API1_R_AG; *<u>MtAPI</u>* <u>region 2</u> (*MtAPI* bases 1549 to 3153), primer API2_F_AG and API2_R_AG; *<u>MtAPI</u>* <u>region 3</u> (*MtAPI* bases 3154 to 3690), primer API3.4.5._F_AG and API1.2.3_R_AG; *<u>MtAPI</u>* <u>region 4</u> (*MtAPI* bases 3691 to 4278), primer API4_HAPI1_5_F_AG and apiMID_hapi1WH2_R_AG; *<u>MtAPI</u>* <u>region 5</u> (*MtAPI* bases 4279 to 4722), primer hapi1MID_apiWH2_F_AG and apiWH2_pl_R_AG; *<u>MtHAPI1</u>* <u>region 1</u> (*MtHAPI1* bases 1 to 1557), primer pl_hapi1SHD_F_AG and HAPI1_1_R_AG; *<u>MtHAPI1</u>* <u>region 2</u> (*MtHAPI1* bases 1558 to 3054), primer HAPI1_2_F_AG and HAPI1_2_R_AG; *<u>MtHAPI1</u>* <u>region 3</u> (*MtHAPI1* bases 3055 to 3594), primer HAPI1_3.4.5._F_AG and HAPI1_1.2.3._R_AG; *<u>MtHAPI1</u>* <u>region 4</u> (*MtHAPI1* bases 3595 to 4044), primer HAPI1_4_API5_F_AG and hapi1MID_apiWH2_R_AG; *<u>MtHAPI1</u>* <u>region 5</u> (*MtHAPI1* bases 4045 to 4488), primers apiMID_hapiWH2_F_AG and hapi1WH2_pl_R_AG.

*MtAPI* and *MtHAPI1* mCitrine-tagged variants were synthesised with flanking Gateway-compatible attL1/2 sites and cloned into pUC57 by Genscript. The *mCitrine* CDS was inserted with a short N-terminal linker sequence (*linker-mCitrine*: GGAGGTGGAGGTGGAGCT) between *MtAPI* CDS bases 1812 and 1813. Four silent mutations introduced single-cut HpAI and BglII restriction sites flanking the insertion site (see Data S1). 14 additional silent base pair mutations were introduced into the *MtAPI* CDS of *API*^*K517R*^*-mCitrine, API*^*K518R*^*-mCitrine, API*^*K520R*^*-mCitrine, API*^*K547R*^*-mCitrine, API*^*3xKtoR*^*-mCitrine* and *API*^*4xKtoR*^*-mCitrine, API*^*ΔA*^*-mCitrine, API*^*ΔB*^*-mCitrine, API*^*ΔC*^*-mCitrine, API*^*ΔAB*^*-mCitrine, API*^*ΔAC*^*-mCitrine, API*^*ΔBC*^*-mCitrine, API*^*ΔABC*^*-mCitrine* and *API*^*GS*^*-mCitrine*, to create single-cut restriction sites flanking: *MtAPI segment A* (BstBI/BsrGI); *segment B* (BspEI/Nhe); *segment C* (SpeI/SacII).

To generate KtoR mutant MtAPI variants, we introduced the following additional mutations: K517R involved switching alanine 1550 and guanine 1551 (AG to GA); K518R involved changing alanine 1553 to guanine (A to G); K520R involved substituting alanine 1559 with guanine (A to G) and K547R involved altering alanine 1640 to guanine (A to G). To generate *API*^*ΔA*^*-mCitrine, API*^*ΔB*^*-mCitrine, API*^*ΔC*^*-mCitrine, API*^*ΔAB*^*-mCitrine, API*^*ΔAC*^*-mCitrine* and *API*^*ΔBC*^*-mCitrine, API*^*ΔABC*^*-mCitrine* and *API*^*GS*^*-mCitrine*, 126bp (*MtAPI segment A*: bases 1527 - 1654), 60bp (*MtAPI segment B*: bases 3789 to 3850) and/or 87bp (*MtAPI segment C*: bases 4035 to 4123) of the *MtAPI* CDS were deleted or swapped with corresponding GS-linker sequences (*GS-linker A*: GGAGGTTCTGGTGGAGGTGGATCAGGTGGAGGATCTGCTGGCTCCGCTGCTGGTTCTGGCGAATTCGGAGGATCTGGAGGTGGAGGATCTGGAGGTGGATCTGCTGGATCTGCGCTGGTTCTGGA; *GS-linker B*: GGATCCGGTGGTGGAGGTTCTGGAGGTTCAGCTGGATCAGCTGCTGGAGGAGGT GGATCC; *GS-linker C*:GGATCAG GAGGTGGAGGTTCTGGAGGTGGATCAGCTGGATCAGCTGCTGGATCAGGTGAATTCGGAGGTTCTGGTGGAGGTGGATCA).

To create mCitrine-tagged *MtHAPI1* variants, the *linker-mCitrine* CDS was inserted between *MtHAPI1* bases 1704 and 1705. Eight silent base pair mutations were introduced into the *MtHAPI1* CDS to generate single-cut restriction sites flanking the *MtAPI segment A* (ClaI /BssHII), *linker-mCitrine* (BspEI/PmlI), *MtAPI segment B* (SpeI/XhoI) and *MtAPI segment C* (BstBI/ MfeI) insertion sites. For generating *HAPI1*^*+ABC*^*-mCitrine* and *HAPI1*^*GS*^*-mCitrine, MtAPI segments* or *GS-linker* sequences were inserted as follows: *MtAPI segment A* or *GS-linker A* (126bp between *MtHAPI1* bases 1536 and 1537), *MtAPI segment B* or *GS-linker B* (60bp, between *MtHAPI1* bases 3690 and 3691), *MtAPI segment C* or *GS-linker C* (87bp between *MtHAPI1* bases 3879 and 3880.

All *42aa-mCitrine* derived constructs feature a variable N-terminal sequence, followed by a *shortAPI-linker-mCitrine* sequence (*MtAPI* bases 1655 to 1812 plus “GGAGGTGGAGGTGGAGCT” linker; in total 177bp) and *mCitrine* with a stop codon. *42aa-mCitrine* was amplified from *pUC57-API-mCitrine* using primers SB295 and SB296. *GS-mCitrine* was amplified from *pUC57-API*^*GS*^*-mCitrine* using primers SB323 and SB296. *41aa-mCitrine* was was amplified from *pKGW-pAtUBQ-42aa-mCitrine* using primers SB329 and SB296. 2*2-43aa-mCitrine* was amplified from *pKGW-pAtUBQ-42aa-mCitrine* using primers SB324 and SB296. *31-43aa-mCitrine* was amplified from *pKGW-pAtUBQ-42aa-mCitrine* using primers SB325 and SB296. *22-30aa-mCitrine* was amplified from *pDONR221-31-43aa-mCitrine* with primers SB343 and SB296. *4xKtoR-mCitrine* was amplified from *pUC57_KAN_API_K4xR_mCitrine* with primers SB342 and SB296. PCR reactions were performed using Phusion DNA polymerase (New England Biolab Inc, UK). The CDSs of *1-21aa-mCitrine, 4xSTtoA-mCitrine, 4xSTtoD-mCitrine*, were synthesised with Gateway-compatible attL1/2 sites and cloned into pUC57 by Genscript. *1-21aa-mCitrine* includes a start codon with *MtAPI* bases 1528 to 1587 fused to the *shortAPI-linker-mCitrine* sequence. *4xSTtoA-mCitrine* and *4xSTtoD-mCitrine* include a start codon with *MtAPI segment A* (bases 1527 to 1654) fused to the *shortAPI-linker-mCitrine* sequence. *4xSTtoA-mCitrine* features the following mutations: AGCA to GCAG (MtAPI bases 1588 to 1591), alanine 1606 and thymine 1612 to guanines, and thymine 1638 to cytosine. *4xSTtoD-mCitrine* features the following mutations: AGCAC to GATGA (MtAPI bases 1588 to 1592), ACA to GAT (MtAPI bases 1606 to 1608), TCA to GAT (MtAPI bases 1612 to 1614).

All entry and destination vectors were validated using diagnostic restriction digest and Sanger sequencing (Source BioScience). Destination clones were assembled by recombining pUC57 gateway compatible *MtAPI* and *MtHAPI1* variants with *pENTR4_1_prAtUBQ3* or *pENTR4_1_prMtAPI* and *pENTR_p2rp3_T35STerm* into *pKGW-RR-MGW* destination vector with LR Clonase Plus (Thermo Fisher Scientific) (*15*). All primer sequences and constructs used in this study are detailed in Table S1 and S2, respectively. A fasta file containing the vector and coding sequences is available as Data S1.

### Generation of transgenic plants

#### Arabidopsis thaliana stable transformations

Destination vectors were introduced into *Agrobacterium tumefaciens* strain *GV3101* (resistant to Tetracycline, Rifampicin, and Gentamicin) and transformed into *A. thaliana* accession *Col-0* and *dis3-4*. Primary transformants were screened for DsRed fluorescence at the seed and seedlings stage (5 days of growth on soil). Screening was performed using a portable NIGHTSEA model SFA with the GR-Green wavelength set (excitation: 510-540nm; emission: 600nm longpass). At least three independent transgenic lines were propagated for each construct and background.

#### Medicago truncatula root transformations

Destination vectors were introduced into *Agrobacterium rhizogenes* strain *Arqua1193* (resistant to Rifampicin and Carbenicillin), and transformed into *A17* and *api* seedling roots according to Limpens et al. (*26*). Transformed roots were subsequently grown on Fahraeus medium for three weeks.

#### Nicotiana benthamiana leaf infiltrations

To transiently express proteins, destination vectors were introduced into *Agrobacterium tumefaciens* strain *GV3101* (resistant to Tetracycline, Rifampicin, and Gentamicin). Overnight bacterial cultures were resuspended in agroinfiltration medium (10mM MgCl_2_, 10mM 2-(N-morpholino)ethanesulfonic acid (MES) pH 5.7 and 200 μM acetosyringone) to an OD of 0.6. Bacterial suspensions were injected into the abaxial side of 4-week-old *N. benthamiana* leaves. Expression of constructs was analysed 3 days post infiltration.

### Bioinformatic analysis of protein sequence conservation and intrinsic disorder *Mt*API/HAPI1 amino acid conservation barcode

To assess amino acid conservation, the protein sequences of *Mt*API and *Mt*HAPI1 were aligned using the EMBOSS Needle pairwise alignment tool (https://www.ebi.ac.uk/jdispatcher/psa/emboss_needle). Alignment results, reflecting the degree of similarity or mismatch between residues in the form of pipe/colon/period and space symbols, were translated into colour-coded stripes using Rstudio. The colour-code was as follows: black (pipe = identical residue), dark grey (colon = highly similar aa), light grey (period = moderately similar aa) and white (space = alignment gap). The percentage of identical residues in each region using NCBI BLASTp search (https://blast.ncbi.nlm.nih.gov/Blast.cgi).

### Intrinsic Disorder

Intrinsically disordered protein regions (IUPRED3) and disordered protein binding regions (ANCHOR2) were predicted using the web server IUPRED3 (https://iupred3.elte.hu/). Sequences were input into IUPRED3 to obtain disorder scores, which were then plotted as line graphs using Rstudio.

### Legume *Mt*API/HAPI like protein alignments

Legume *Mt*API and *Mt*HAPI1-like sequences were aligned using the “Create Alignment” function in CLC Main Workbench 20 with settings of gap open cost 10, gap extension cost 1, and the “very accurate” alignment algorithm. The resulting alignment graphics were imported into Inkscape, where text styles were adjusted to match the overall figure design.

### Phylogenetic Analysis

Legume SCAR protein sequences were retrieved from publicly available legume reference genomes using a BLASTp search with the *At*SCAR2 sequence (*At2G38440*; Tair) as query. Sequences for *Lupinus angustifolius, Glycine max, Phaseolus vulgaris, Vigna angularis* and *Trifolium pratense* were obtained from the EnsemblPlants platform (http://plants.ensembl.org/index.html); for *Nissolia schottii, Arachis hypogaea, Cajanus cajan* and *Lablab purpureus* from the SymDB database (https://www.polebio.lrsv.ups-tlse.fr/symdb/web/); for *Lotus japonicus MG20* from Lotus Base (https://lotus.au.dk/); and for *Medicago truncatula* from the Medicago A17 genome browser (https://medicago.toulouse.inra.fr/MtrunA17r5.0-ANR/). Full-length sequences were aligned and subsequently automatically trimmed using the bioconda packages “mafft” (version 7.471) (https://anaconda.org/bioconda/mafft) (*27*) and “trimAI” (version 1.4.1) (https://trimal.readthedocs.io/en/latest/) (*28*). Phylogenetic trees were constructed using the bioconda package “IQ-TREE” (version 2.0.3) (*29*) with maximum likelihood and bootstrapping (1000Ufboot+SH-aLRT) and identified the best fitting substitution models: “JTT+G4” for Fig. 1A and “HIVb+F+I+G4” for fig. S3A. Trees were visualised with the online tool “interactive Tree Of Life” (iTOL version 6.9.1 :https://itol.embl.de/) (*30*) and stylistically modified for figure presentation using Inkscape.

### Microscopic analysis of root hair and trichome morphology

#### M. truncatula root hair length analysis

Three weeks post-transformation, epifluorescence microscopy images of DsRed-expressing *M. truncatula* roots were captured using a Leica M165 FC Fluorescent Stereomicroscope with a DFC310FX camera and DSR filter (10447412). All images were taken at the same magnification (including a reference scale) with the Leica Application Suite Software (Version 4.8.0). Root hair length was quantified using the ROI manager and Freehand line tools in ImageJ2. The image scales were globally calibrated using the SetScale function. For each genotype, the length of 20 root hairs from 5 independently transformed roots (n=100) was measured in millimetres. Data normality was assessed using the Shapiro-Wilk test, and for p-values below 0.5, the Kruskal-Wallis test with Bonferroni p-value adjustment (alpha = 0.05) was applied for posthoc analysis. Box plots were generated in R. Images were adjusted for greyscale, brightness, and contrast using GIMP, and figure panels were assembled in Inkscape with manually added scale bars based on the reference scale.

#### A. thaliana trichome branch length analysis

Scanning electron micrographs (SEM) of *A. thaliana* leaves and trichomes were captured using the Hitachi tabletop microscope TM4000 Plus with the provided software. Leaves of nine-day-old *A. thaliana* plants, grown on soil, were detached and mounted on sample stubs with water droplets. The cooling stage was set to −25°C with low vacuum applied upon sample insertion. BSE (backscatter) detector mode was used with an accelerating voltage of 15 kV. Whole leaf images were captured at 30x magnification, and trichome close-ups at 150x magnification, with corresponding scale bars calculated by the software. Brightness, contrast and cropping adjustments were made using GIMP. Three images from independent transgenic lines per genotype were analysed. For each image, five representative trichomes were selected, and the length of each trichome branch was measured from branch points to tip using the Freehand Line tool in ImageJ2. The shortest branch ratio was determined by dividing the length of the shortest branch by the longest branch for each trichome (1 = equal length; n =15). This data was filtered, analysed and visualised as box plots In R. Final figure panels, including representative trichome images, scale bars, and barplots were assembled using Inkscape.

### Localisation and expression analysis via confocal microscopy

For confocal imaging, of *A. thaliana* root cells and *N. benthamiana* pavement cells, samples were mounted in water on glass slides (Fisher, 1-1.2mm) and covered with coverslips (Epredia, #1.5). Imaging was performed using a LeicaTCS SP8 upright confocal microscope, equipped with a white light laser system.

### Imaging of *A. thaliana* seedling roots

Five-day-old stably transformed *A. thaliana* seedlings, grown on ½ MS pH 5.7, were mounted and screened for root fluorescence. Single-plain root cell images were acquired with HC PL APO CS2 63w/1.20 water immersion objective and the following laser/detector settings: Sequential scan; sequence1 (mCitrine): excitation with WLL line 518nm: 70% intensity, detection with HyD1: 523-535nm gain [%] 500, time gating on, 0.3-6ns; sequence 2 (DsRed): excitation WLL line 570nm: 5% intensity, detection with PMT3 577-602nm gain [V] 890.8 offset: −0.02. Unidirectional scan, speed: 200 Hz, line averaging: 8, pinhole: 1 airy unit, zoom: 3.

### Imaging of *N. benthamiana* pavement cells

Three days after transformation, leaf discs (cork borer size three) of six independently transformed plants were analysed for fluorescence. Single-plane, 16-bit images of the pavement cell were acquired with the following settings: Sequential scan; sequence 1 (mCitrine): excitation 515nm: 0.5%-10% intensity, detection with HyD1: 518-537nm gain [%] 500, time gating 0.6 - 6ns; sequence 2 (DsRed): excitation 560nm: 0.5%-10% intensity, detection with HyD5 574-598nm gain [%] 500, time gating 0.4 - 6ns. Unidirectional scan, speed: 600 Hz, line averaging: 8, pinhole: 1 airy unit, zoom: 1. Laser intensities were optimised between experiments while maintaining consistent settings within each experiment.

To calculate mCitrine/dsRed ratios, the segment and ratio tool of the FRETENATOR2 beta version (https://github.com/JimageJ/FRETENATOR2; based on Rowe et al., 2023 (*31*)) FIJI plug-in was used. The plug-in automatically segments images and calculates ratios for the segmented pixels areas. Setting were chosen as follows: Denominator (‘Donor’) and Segmentation channel: DsRed; Numerator (‘Acceptor’) channel: mCitrine; Gaussian segmentation, maximum intensity: 65534 without background subtraction; pixel by pixel analysis: on. For each genotype, mean ratio measurements were taken from one image per six independently transformed plants, using the ‘Maximum z of emission ratio z projection X1000’ images (n = 6). The mean ratios were divided by 1000 and plotted using R.

### Quantitative reverse transcription PCR (qRT-PCR) analysis

For each construct, three leaf discs (cork borer size five) were collected from a single transformed *N. benthamiana* leaf and immediately frozen in liquid nitrogen. As negative control, three leaf discs per biological replicate were also harvested from untransformed leafs. A total of six biological replicates per genotype was analysed. RNA extraction was carried out following the manufacturer’s protocol for the RNEasy Plant Mini Kit (QIAGEN), using buffer RLC. For each sample, 4µg of total RNA was used as a template for cDNA synthesis, which was performed using the Transcriptor First Strand cDNA Synthesis Kit (Roche). Quantitative reverse transcription PCR (qRT-PCR) was conducted using 2.5 μl of a 1:8 dilution of the first-strand cDNA and LightCycler 480 SYBR Green I Master mix, following the manufacturer’s instructions (Roche). *mCitrine* transcript levels were analysed using primers SB268 and SB269 (Table S1). The *N. benthamiana* genes *NtL23* (*Niben101Scf01444g02009*) and *NtFBOX* (*Niben101Scf04495g02005*) were chosen as constitutively expressed reference genes (*32*). *NtL23* was amplified using primers SB304 and SB305 (Table S1). *NtFBOX* was amplified using primers SB306 and SB307 (Table S1). The expression of *mCitrine* was normalised to *NtL23* an *NtFBOX* expression using the efficiency-corrected ΔΔCq method. The data was visualised using R software.

### SDS-PAGE and Immunoblot

For each construct, two leaf discs (cork borer size five) were collected from six independently transformed *N. benthamiana* plants, pooled, and immediately frozen in liquid nitrogen. The frozen tissue was ground into a fine powder with a porcelain pestle and mortar. To this powder, 500µl of lysis buffer (GTEN: 50mM Tris-HCl pH7.5, 10% [vol/vol] glycerol, 1mM EDTA, 150mM NaCl; supplemented with 2% [wt/vol] polyvinylpolypyrrolidone, 0.1% [vol/vol] Tween-20, phosphatase inhibitors (Sigma P5726 and P0044) and plant protease inhibitors (Sigma P9599)) was added. The mixture was incubated on ice for 10 minutes. Following a 5-minute centrifugation at maximum speed (4°C), 400µl of the supernatant was mixed with 4x Lämmli buffer containing beta-mercaptoethanol and boiled at 95°C for five minutes.

SDS-polyacrylamide gel electrophoresis (PAGE) was carried out using 4-20% Mini-PROTEAN TGX Precast protein gels (Bio-Rad). After electrophoresis, proteins were transferred onto a 0.45µm PVDF membrane (Immobilon, Merck). The membranes were stained with a 1:1 mix of glacial acetic acid and Ponceau Red (Bio-Rad) for protein visualisation, followed by three brief washes in deionized water. Membranes were blocked for 1 hour in a TBS-T buffer (20mM Tris Base, 150mM NaCl, 0.1% [vol/vol] Tween20) containing 5% [wt/vol] milk powder. Subsequent incubations (for 1 hour) were carried out in primary and secondary antibodies diluted in the same blocking solution. After each incubation step, membranes were washed three times for 10 minutes with TBS-T buffer. The following antibodies and dilutions were used: mouse anti-GFP (B2 sc-9996, Santa Cruz biotechnology) 1:2500, rabbit anti-RFP (AB62341, Abcam), m-IgG Fc binding protein HRP (sc-542732, Santa Cruz biotechnology) 1:5000 and goat anti-rabbit IgG HRP (AB205718, Abcam) 1:5000. Protein bands were detected using chemiluminescence with ECL Plus (Thermo Fisher Scientific) and visualised on an Amersham Imager 600.

### Protein extraction, immunoprecipitation and Mass spectrometry analysis

Three leaves from *N. benthamiana* plants, transformed with either *mCitrine* or *42aa-mCitrine*, were harvested 72 hours post-agroinfiltration. The leaves were flash-frozen in liquid nitrogen and ground into a fine powder using porcelain pestles and mortars. Proteins were extracted by adding 5 mL of lysis buffer (GTEN: 50 mM Tris-HCl, pH 7.5, 10% [vol/vol] glycerol, 1 mM EDTA, 150 mM NaCl, supplemented with 2% [wt/vol] polyvinylpolypyrrolidone, 0.1% [vol/vol] Tween-20, phosphatase inhibitors [Sigma P5726 and P0044], and plant protease inhibitors [Sigma P9599]) to the leaf powder. For immunoprecipitation, 60 µL of GFP-Trap® agarose bead slurry (Chromotek) was added to each sample after centrifugation at maximum speed for 5 minutes at 4°C. The samples were incubated for 2 hours at 4°C with slow rotation. Following incubation, the samples were centrifuged at 2000 rpm for 2 minutes at 4°C, and the resulting pellet was washed five times with 1 mL of immunoprecipitation (IP) buffer (GTEN, supplemented with 0.1% [vol/vol] Tween-20). After the final wash, the supernatant was removed, and 20 µL of IP buffer along with 10 µL of 4x Lämmli buffer (supplemented with beta-mercaptoethanol) was added. The sample proteins were denatured by heat treatment at 95°C for 5 minutes. SDS-polyacrylamide gel electrophoresis (PAGE) was performed using 4–20% Mini-PROTEAN TGX Precast protein gels (Bio-Rad). After electrophoresis, the gel was stained overnight in 20 mL SimplyBlue™ (Invitrogen) staining solution, supplemented with 2 mL of 20% NaCl in water (w/v). Protein bands, corresponding to mCitrine and 42aa-mCitrine, were excised and subjected to in-gel reduction and alkylation, followed by trypsin digestion and peptide extraction by the Cambridge Proteomics Centre. The resulting peptides were analysed by liquid chromatography-tandem mass spectrometry (LC-MS/MS) and post translational modification identification was performed using the MASCOT search algorithm with *N. benthamiana* v1.01 gene models (https://solgenomics.net). Additionally, *Mt*API segment A (*Mt*API amino acids 510 - 551) was also analysed for potential post-translational modification sites with the web server application MusiteDeep (https://www.musite.net/), which uses a deep-learning framework for protein post-translational modification site prediction.

### Software versions

The following software versions were used: RStudio 2022.02.1 with R 4.1.3.; Inkscape 1.2.2; ImageJ2 software version 2.14.0 including FIJI plugins; GNU image manipulation program (GIMP version 2.10.30).

### Accession numbers/identifiers

MtAPI (Medtr4g013235/MtrunA17_Chr4g0004861)

MtHAPI1 (Medtr7g071440/MtrunA17_Chr7g0244031)

AtSCAR2/DIS3 (AT2G38440)

AtSCAR1 (AT2G34150), AtSCAR3 (AT1G29170), AtSCAR4 (AT5G01730)

AtNAP1 (AT2G35110)

NtL23 (Niben101Scf01444g02009)

NtFBOX (Niben101Scf04495g02005)

## Acknowledgements

We thank Daniel Szymanski for providing us with A. thaliana Col and dis3-4 seeds, Jim Rowe for help with the installation and usage of the Fretenator plugin, Radoslaw Kowalczyk for technical assistance. We are also grateful to Philip Carella and Alan Wanke for discussion and critical reading of the manuscript.

## Author contributions

Conceptualization: S.B., A.G. and S.S.; Methodology and Investigation: S.B., A.G., M.M., G.C. and A.U.; Supervision: S.B., A.G. and S.S.; Visualisation: S.B.; Writing-original draft: S.B., A.G. and S.S.; Funding acquisition: S.S.

## Declaration of generative AI and AI-assisted technologies in the writing process

The authors used ChatGPT to check grammar and phrasing of the manuscript. After using ChatGPT, the authors reviewed and edited the content as needed and take full responsibility for the content of the publication

## Data and materials availability

All data needed to evaluate the conclusions in the paper are present in the paper and/or the Supplementary Materials. Materials are available on request from S.S.

