## Supplemental Figures and Tables for "A molecular basis for plant SCAR/WAVE functional divergence"

Sabine Brumm et al.

*Sebastian Schornack.

**This PDF files includes:**

Figs. S1 to S8

Table S1 and S2

**Other Supplementary Materials for this manuscript include the following:**

Data S1

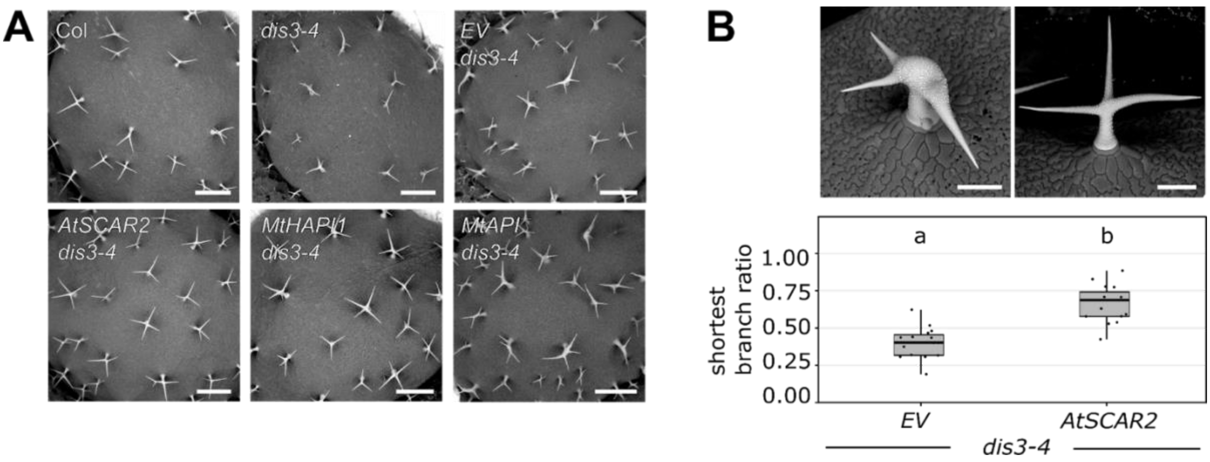


**Fig. S1: The *M. truncatula SCAR* gene *HAPI1* rescues the *A. thaliana dis3-4* trichome development phenotype to similar levels as At*SCAR2*.**

**(A)** Scanning electron micrographs of *A. thaliana* leaves from untransformed Col, *dis3-4* lines and *dis3-4* lines transformed with *empty vector* (EV), *AtSCAR2*, *MtHAPI1* or *MtAPI* driven by the *AtUBQ3* promoter. Scale bars, 0.5 mm. (**B)** Controls for the trichome branch length rescue in *A. thaliana dis3-4* (Fig1): *EV* (negative) and *AtSCAR2* (positive). Scale bars, 90 µm. Shortest branch ratio (n = 15 / genotype). Statistics: Shapiro-Wilk test, followed by Kruskal-Wallis with Bonferroni p-value adjustment (alpha = 0.05); significance differences are indicated by letters a and b.

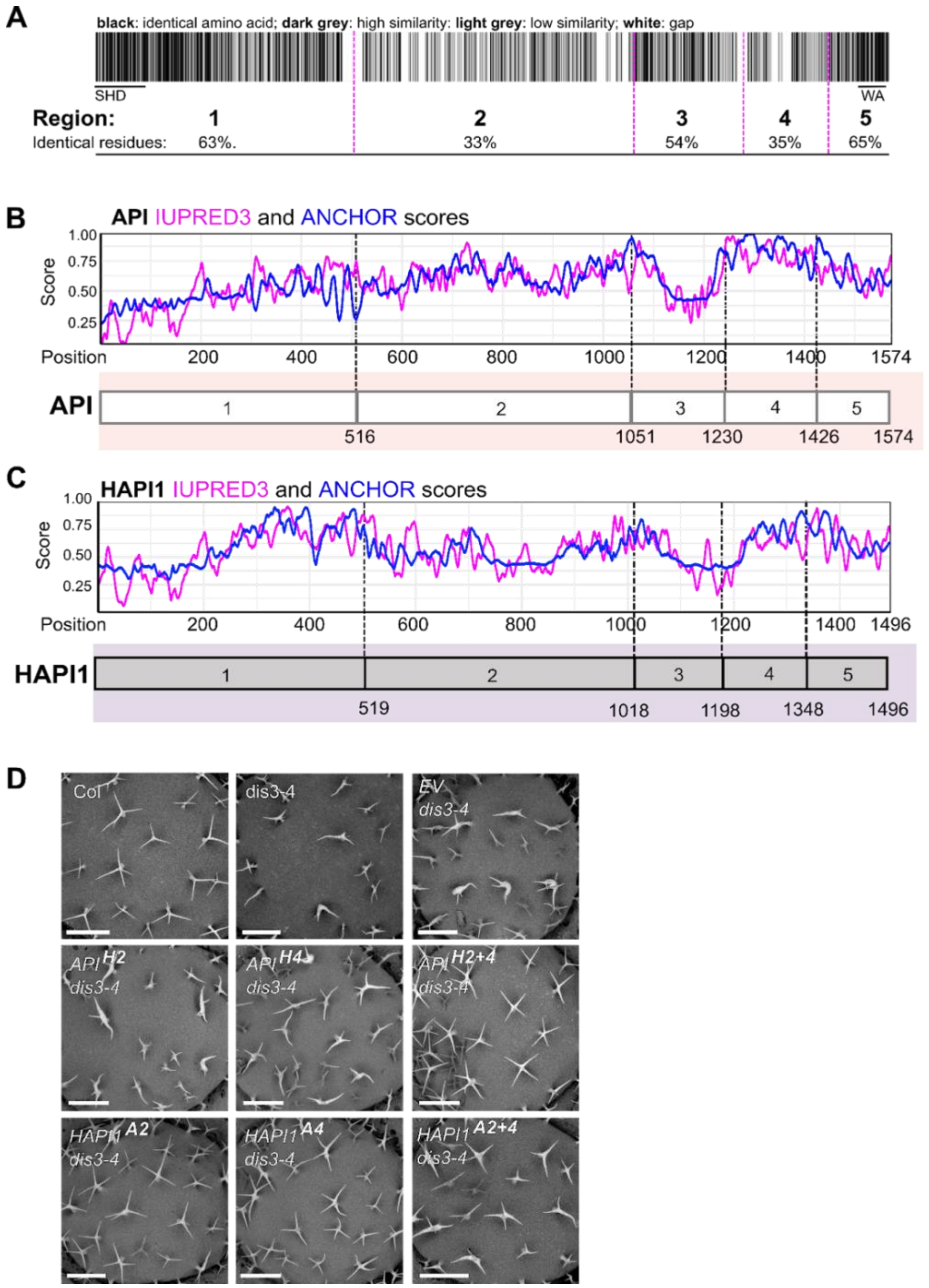


**Fig. S2: Bioinformatic analysis of *Mt*API and *Mt*HAPI1 protein sequences and *A. thaliana* complementation studies with chimeric proteins.**

**(A)** Schematic barcode representation of *Mt*API and *Mt*HAPI1 amino acid conservation. Colours represent identical (black), highly similar (dark grey), moderately similar (light grey) amino acid residues, and alignment gaps (white). (**B** and **C)** Prediction of intrinsically disordered protein regions (IUPRED3, pink) and disordered protein binding regions (ANCHOR2, blue) by IUPRED3 in *Mt*API (B) and *Mt*HAPI1 (C). **(D)** Scanning electron micrographs of *A. thaliana* leaves from Col, *dis3-4*, and *dis3-4* lines transformed with *empty vector* (EV) and chimeric *API/HAPI1* variants under the *AtUBQ3* promoter. Scale bars, 0.5 mm.

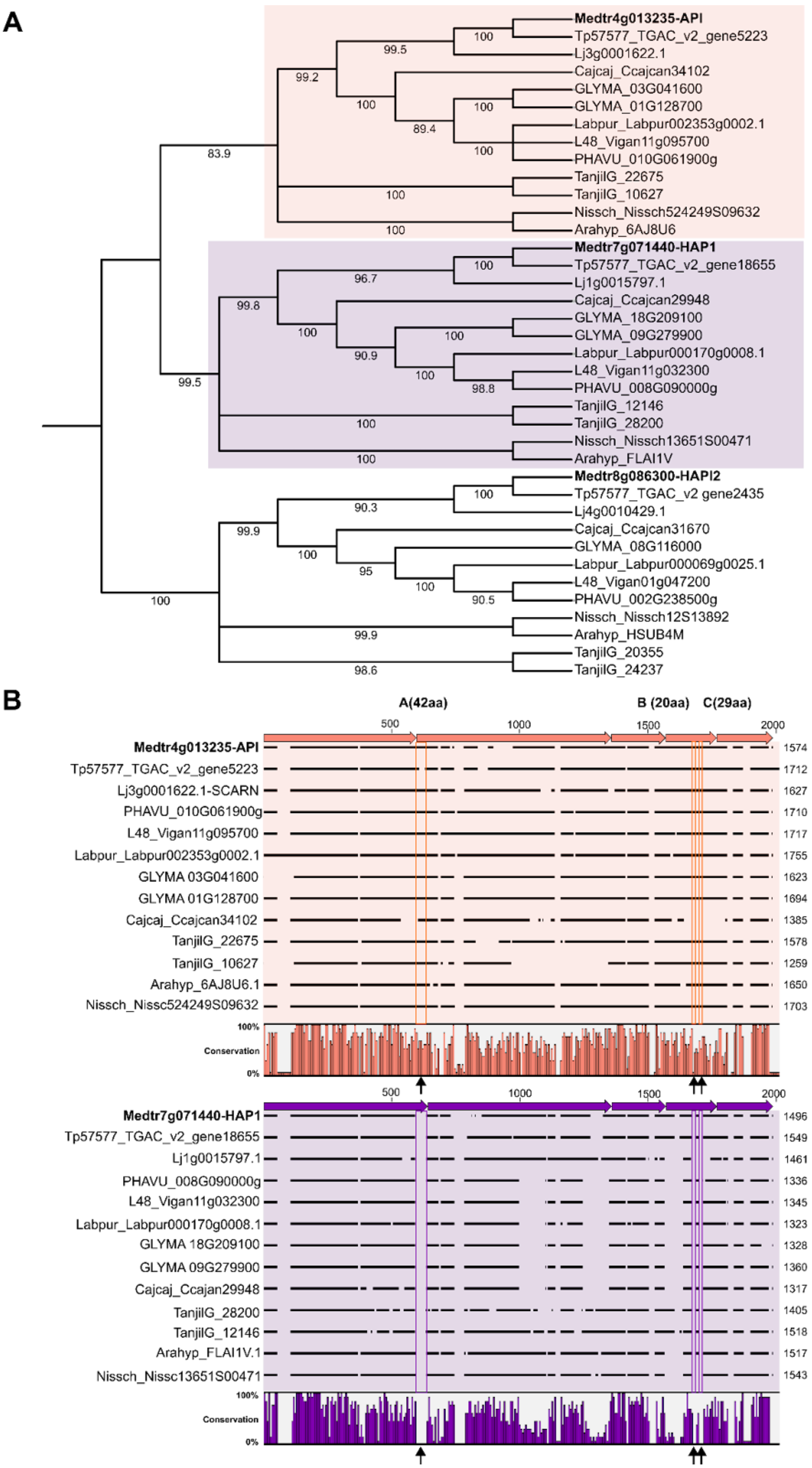
A
Medtr4g013235-API
Tp57577_TGAC_v2_gene5223
Lj3g0001622.1
Cajcaj_Ccajcan34102
GLYMA_03G041600
GLYMA_01G128700
Labpur_Labpur002353g0002.1
L48_Vigan11g095700
PHAVU_010G061900g
TanjilG_22675
TanjilG_10627
Nissch_Nissch524249S09632
Arahyp_6AJ8U6
Medtr7g071440-HAP1
Tp57577_TGAC_v2_gene18655
Lj1g0015797.1
Cajcaj_Ccajcan29948
GLYMA_18G209100
GLYMA_09G279900
Labpur_Labpur000170g0008.1
L48_Vigan11g032300
PHAVU_008G090000g
TanjilG_12146
TanjilG_28200
Nissch_Nissch13651S00471
Arahyp_FLAI1V
Medtr8g086300-HAPI2
Tp57577_TGAC_v2 gene2435
Lj4g0010429.1
Cajcaj_Ccajcan31670
GLYMA_08G116000
Labpur_Labpur000069g0025.1
L48_Vigan01g047200
PHAVU_002G238500g
Nissch_Nissch12S13892
Arahyp_HSUB4M
TanjilG_20355
TanjilG_24237
B
A(42aa)
B (20aa)
C(29aa)
500
1000
1500
2000
Medtr4g013235-API 1574
Tp57577_TGAC_v2_gene5223 1712
Lj3g0001622.1-SCARN 1627
PHAVU_010G061900g 1710
L48_Vigan11g095700 1717
Labpur_Labpur002353g0002.1 1755
GLYMA 03G041600 1623
GLYMA 01G128700 1694
Cajcaj_Ccajcan34102 1385
TanjilG_22675 1578
TanjilG_10627 1259
Arahyp_6AJ8U6.1 1650
Nissch_Nissc524249S09632 1703
100%
Conservation
0%
Medtr7g071440-HAP1 1496
Tp57577_TGAC_v2_gene18655 1549
Lj1g0015797.1 1461
PHAVU_008G090000g 1336
L48_Vigan11g032300 1345
Labpur_Labpur000170g0008.1 1323
GLYMA 18G209100 1328
GLYMA 09G279900 1360
Cajcaj_Ccajan29948 1317
TanjilG_28200 1405
TanjilG_12146 1518
Arahyp_FLAI1V.1 1517
Nissch_Nissc13651S00471 1543

**Fig. S3: Sequence conservation analysis of *Mt*API and *Mt*HAPI1.**

**(A)** Maximum likelihood phylogenetic tree of SCAR proteins from various legume species, showing three distinct clades. *Mt*API (orange) and *Mt*HAPI1 (purple) like proteins are more closely related to each other than to *Mt*HAPI2-like proteins. Node bootstrap values are indicated. Species abbreviations: Medtr (*Medicago truncatula*), Lj (*Lotus japonicus MG20*), TanjilG (*Lupinus angustifolius*), LR48_Vigan (*Vigna angularis*), PHAVU (*Phaseolus vulgaris*), Tp (*Trifolium pratense*), GLYMA (*Glycine max*), Nissch (*Nissolia schottii*), Arahyp_arahy (*Arachis hypogaea*), Cajcaj_Ccajan (*Cajanus cajan*), Labpur (*Lablab purpureus*). **(B)** Sequence conservation analysis of *Mt*API and *Mt*HAPI1-like legume proteins identifies three unique amino acid segments in *Mt*API-like sequences (orange boxes over black arrowheads) absent in *Mt*HAPI1-like sequences (purple boxes over black arrowheads). The five regions of varying conservation are indicated by orange and purple arrows. Black lines represent aligned amino acids, while white spaces indicate alignment gaps. The total numbers of amino acids per sequence are shown at the end of each alignment. Histograms below the alignments show the percentage of conservation for each position (orange bars for *Mt*API-like, purple bars for *Mt*HAPI1-like sequences).

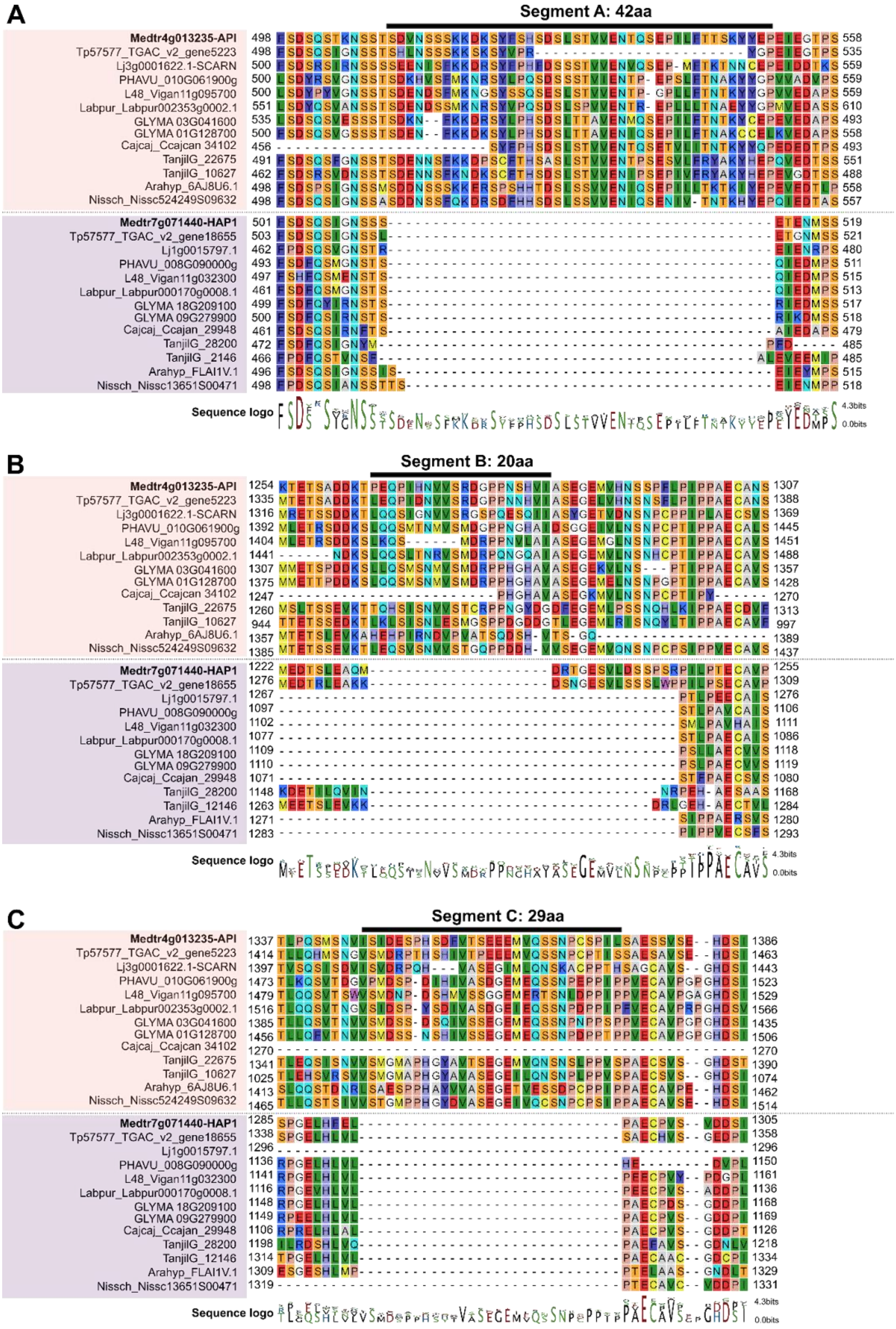
A
Segment A: 42aa
Medtr4g013235-API
Tp57577_TGAC_v2_gene5223
Lj3g0001622.1-SCARN
PHAVU_010G061900g
L48_Vigan11g095700
Labpur_Labpur002353g0002.1
GLYMA 03G041600
GLYMA 01G128700
Cajcaj_Ccajcan 34102
TanjilG_22675
TanjilG_10627
Arahyp_6AJ8U6.1
Nissch_Nissc524249S09632
Medtr7g071440-HAP1
Tp57577_TGAC_v2_gene18655
Lj1g0015797.1
PHAVU_008G090000g
L48_Vigan11g032300
Labpur_Labpur000170g0008.1
GLYMA 18G209100
GLYMA 09G279900
Cajcaj_Ccajan_29948
TanjilG_28200
TanjilG_2146
Arahyp_FLAI1V.1
Nissch_Nissc13651S00471
Sequence logo
4.3bits
0.0bits
B
Segment B: 20aa
Sequence logo
C
Segment C: 29aa
Sequence logo

**Fig. S4: Detailed alignment of *Mt*API segments A, B, and C in closely related legume homologs.**

(**A** to **C**) Detailed sections of the alignment from fig. S3, focusing on *Mt*API segments A (A), B (B), and C (C). Amino acid positions at the start and end of each alignment are indicated. The backgrounds of amino acids are coloured according to the RasMol colour scheme. Orange: *Mt*API-like sequences; Purple: *Mt*HAPI1-like sequences. Segments A, B, and C are highlighted by black lines above each alignment.

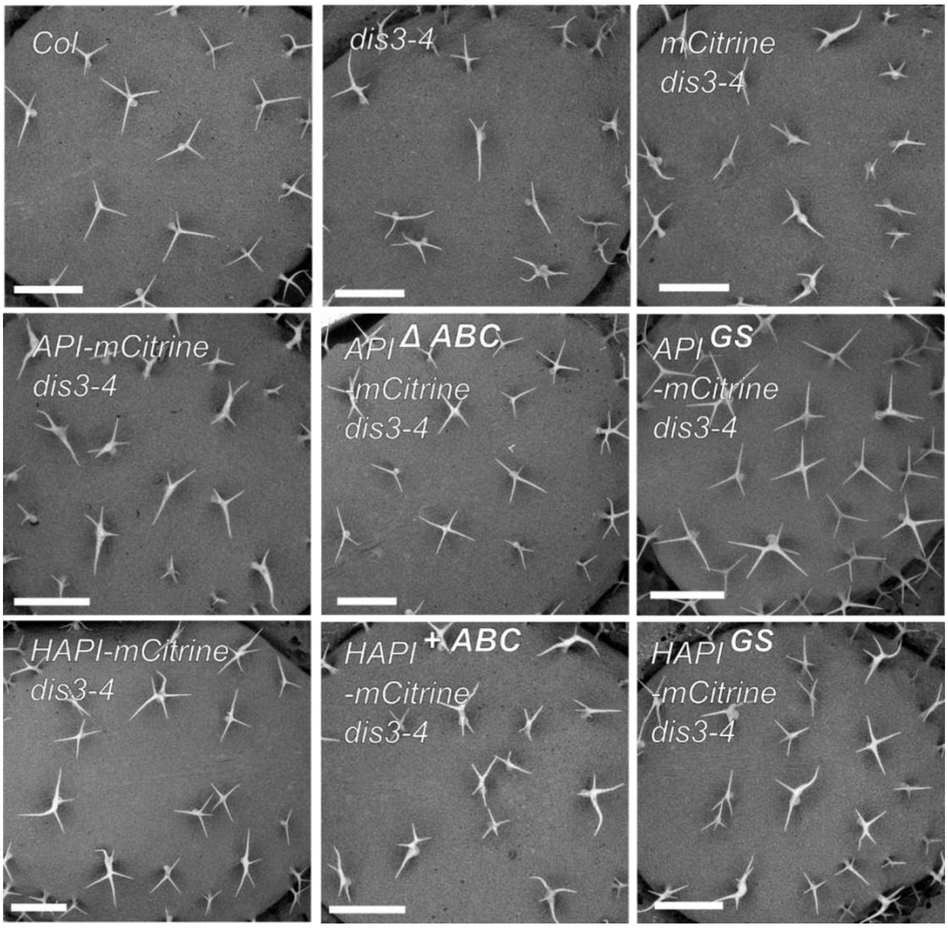


**Fig. S5: *A. thaliana* leaf overview from complementation studies with MtAPI and MtHAPI1 segment A, B, C, and GS-linker variants.**

Scanning electron micrographs of *A. thaliana* leaves from Col, *dis3-4*, and *dis3-4* lines expressing either an empty vector (EV) or mutant *MtAPI* or *MtHAPI1* variants (segments A, B, C, and GS-linkers) under the *AtUBQ3* promoter. Scale bars, 0.5 mm.

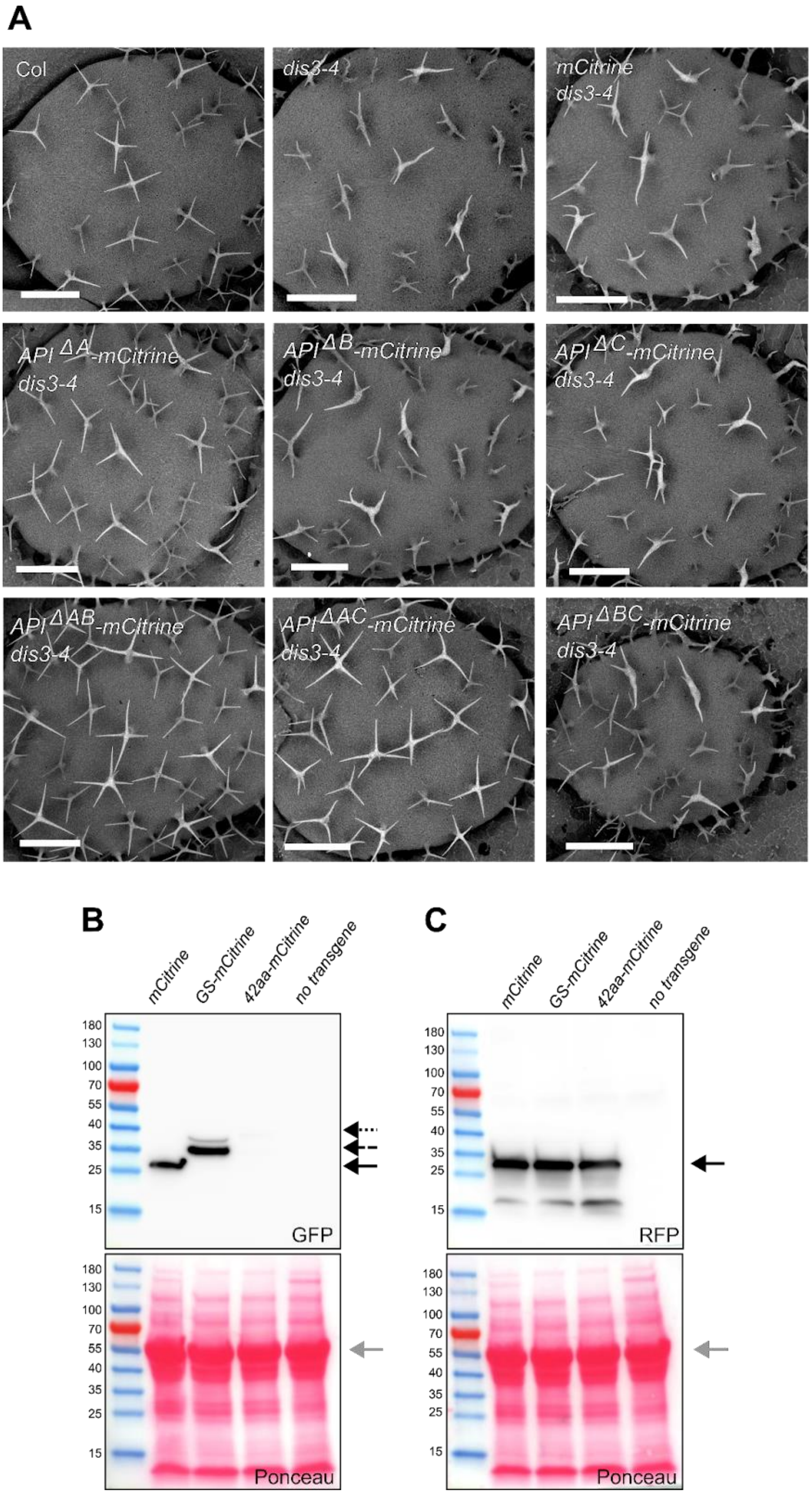


**Fig. S6: Impact of segment A on *Mt*API function in *A. thaliana* trichome development.**
(**A)** Scanning electron micrographs of *A. thaliana* leaves from Col, *dis3-4*, and *dis3-4* lines either expressing empty vector (EV) or *Mt*API segment A, B and C deletion variants under the

*AtUBQ3* promoter. Scale bars, 0.5 mm. (**B** and **C)** Western Blot analysis of mCitrine, GS-mCitrine, 42aa-mCitrine, API-mCitrine and coexpressed dsRed in transiently transformed *N. benthamiana* leaves, with dsRed serving as a positive control. Samples were run on two gels, stained with Ponceau, and probed either with GFP (B) or RFP (C) antibodies. Arrows indicate expected sizes of protein band: Rubisco (55kDa, grey), mCitrine (27kDa, black), dsRed (28kDa, black) GS-mCitrine (36kDa, black with long dotted line) and weakly detectable 42aa-mCitrine (38kDa, black with short dotted line). Prestained protein ladder: Page Ruler 10-180kDa.

**Fig. S7.**

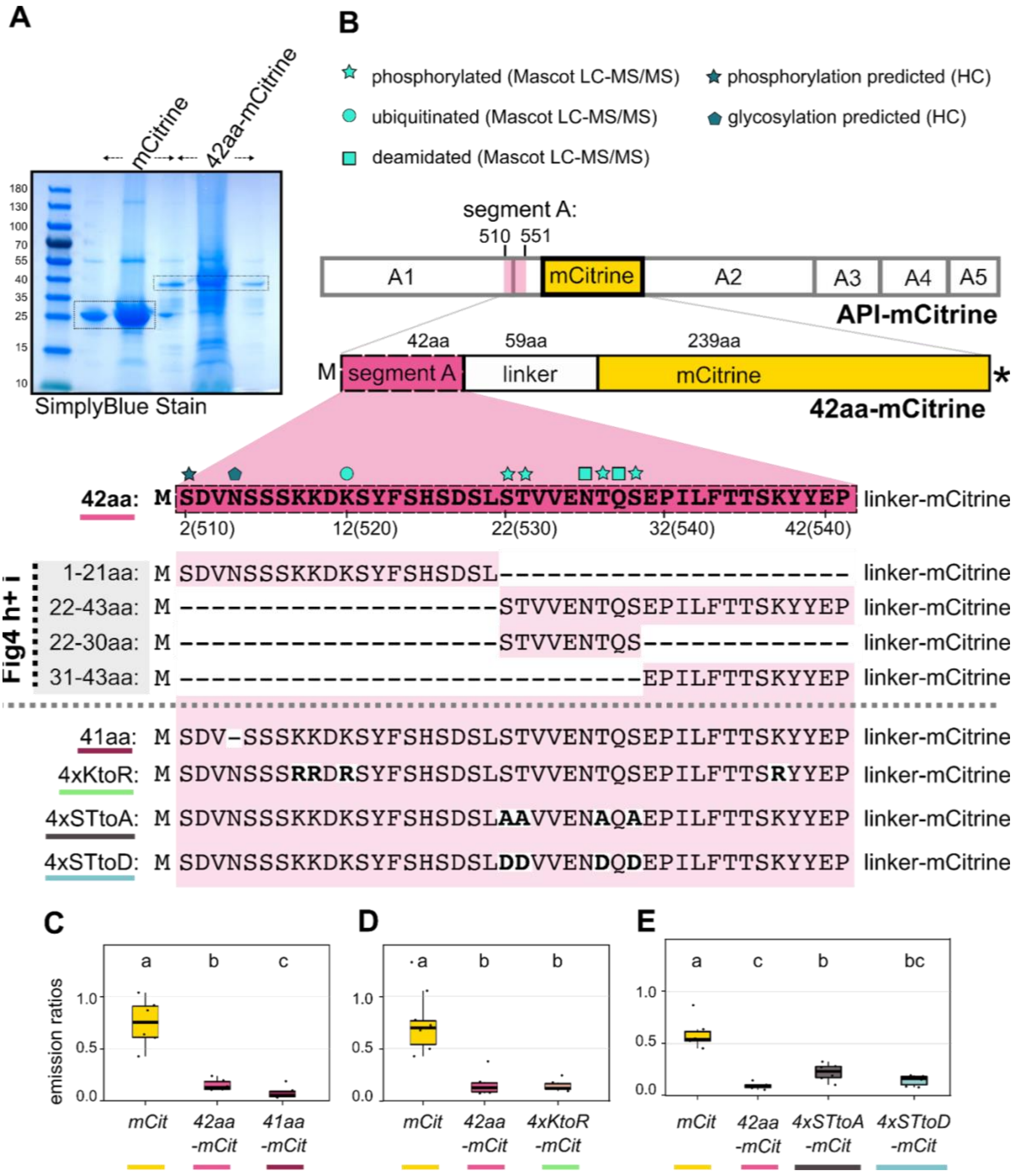


**Fig. S7: Post translational modification analysis of segment A.**

(**A**) SDS gel of immunoprecipitated mCitrine and 42aa-mCitrine proteins stained with SimplyBlue. Arrows indicated spillovers; dotted lines mark gel sections used for LC-MS/MS analysis (**B**) Schematic of API-mCitrine, 42aa-mCitrine, and derived deletion/substitution constructs. Predicted and experimentally validated post- translational modification (PTM) sites in API segment A are mapped onto the 42-amino acid sequence. Numbers indicate amino acid positions in 42aa-mCitrine, with API-mCitrine positions in brackets. PMTs were predicted using MusiteDeep (https://www.musite.net/), with high-confidence (HC, score > 0.5) sites indicated by symbols with dark filling. Experimentally confirmed PTMs (via LC-MS/MS) are marked with cyan symbols. PTM types: phosphorylation (star), ubiquitination (circle), deamination (square), glucosylation (pentagon). (**C** to **E**) Quantification of mCitrine fluorescence in 42aa-mCitrine derived constructs relative to dsRed. 41aa-mCitrine (C), 4xKtoR-mCitrine (D), 4xSTtoA-mCitrine and 4xSTtoD-mCitrine (E) ratios compared to mCitrine (mCit) and 42aa-mCitrine controls. mCitrine/dsRed signals were calculated using the FIJI plugin FRETENATOR (n =6). The different constructs are colour coded. Statistics: Shapiro-Wilk test Kruskal-Wallis with Bonferroni correction; Statistics: Shapiro-Wilk test, followed by Kruskal-Wallis with Bonferroni correction; significance difference groups indicated by letters a, b, c.

**Fig. S8.**

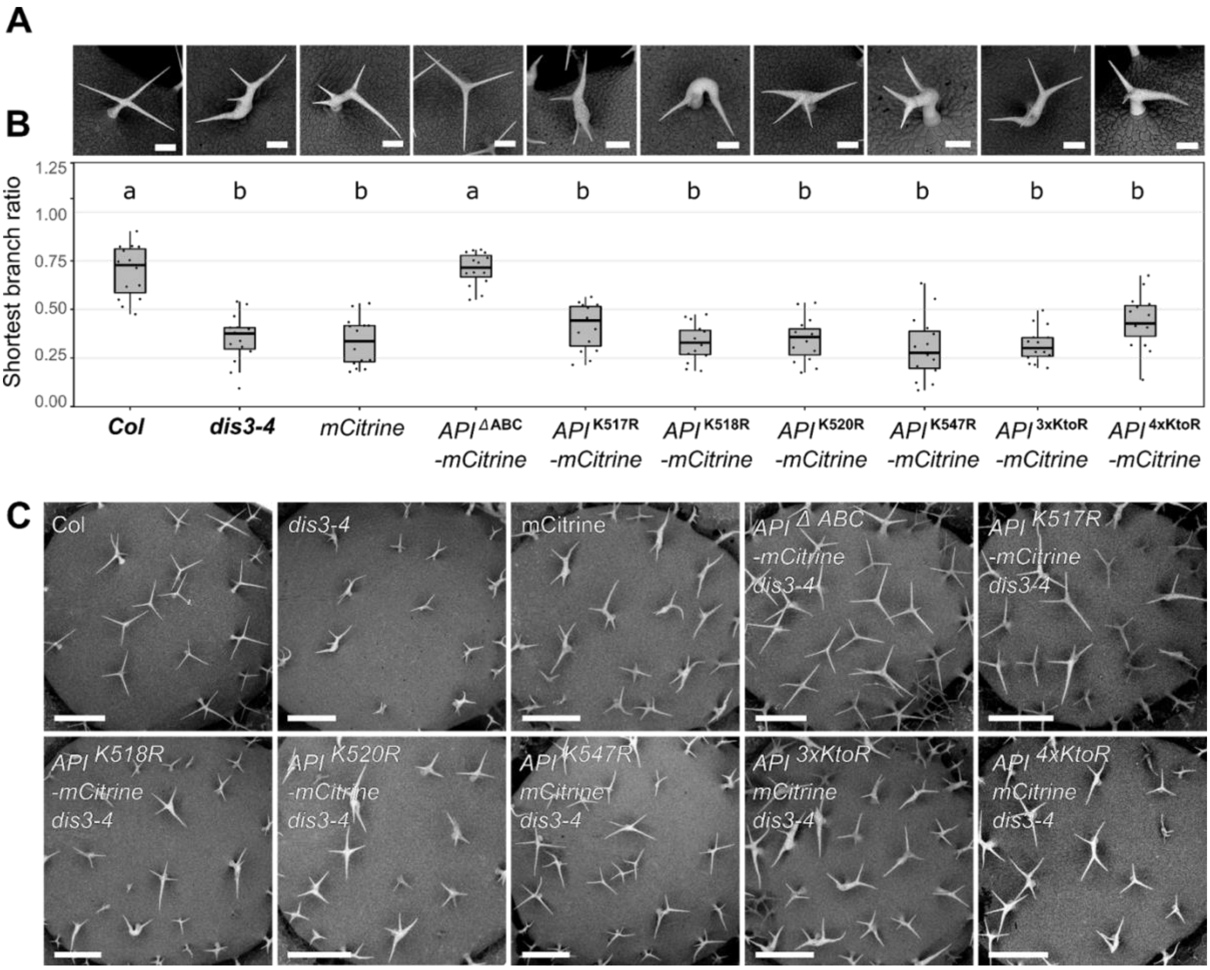


**Fig. S8: Expression of API-mCitrine KtoR mutants in *A. thaliana*.**
(**A**) Scanning electron micrographs of *A. thaliana* trichomes from Col, *dis3-4*, and *dis3-4* lines expressing mCitrine and *Mt*API segment A lysine substitution variants under the *AtUBQ3* promoter. Scale bars, 90µm. (**B**) Lysine-to-arginine (KtoR) substitutions in *Mt*API segment A do not affect *Mt*API functionality in the *A. thaliana dis3-4* mutant. Shortest trichome branch ratio (n=15/genotype). Statistical significance differences (Shapiro-Wilk test, followed by Kruskal-Wallis with Bonferroni correction) are indicated by letters a and b (**C**) Scanning electron micrographs of *A. thaliana* leaves from trichome magnification in (B). Scale bars, 0.5mm.

**Table S1**

| Primer name | Primer sequence |
|---|---|
| pl_apiSHD_F_AG | gatccaagcttctgcagcccATGCCGATATCGAAGTATC |
| API1_R_AG | tatggggaagctggttGGAAGAACTATTTACATCTGACG |
| API2_F_AG | aaatatgtcatccAAGAAAG ATAAATCTTATTTCTCTCATTC |
| API2_R_AG | gtttcattggatcCACAGCATACTGTGTAGAGTTG |
| API3.4.5._F_AG | aaagcatgtcatgGATCCATTGAAGTCTCTTCTC |
| API1.2.3_R_AG | aaacagcacattc AGCAGGTAGTATTGGTGC |
| API4_HAPI1_5_F_AG | aatactgcctgctGCATATACTCTCTCTGGAGATG |
| apiMID_hapi1WH2_R_AG | ttggtggtgaAATACCCAATTGTCTCTGC |
| hapi1MID_apiWH2_F_AG | ctcacctgagTCGCTCATGGTGCCACCA |
| apiWH2_pl_R_AG | tcgagggtacctctagacccTCAAGAATCACTCCAACTATCCTCATC |
| pl_hapi1SHD_F_AG | gatccaagcttctgcagcccATGCCTATCTCAAGGTAC |
| HAPI1_1_R_AG | atttatctttcttGGATGACATATTTTCAGTCTC |
| HAPI1_2_F_AG | aaatagttcttccAACCAGCTTCCCCATACTG |
| HAPI1_2_R_AG | acttcaatggatcCATGACATGCTTTGTTGG |
| HAPI1_3.4.5._F_AG | acagtatgctgtgGATCCAATGAAACCTCTACTTC |
| HAPI1_1.2.3._R_AG | cagagagagtatatgcAGCAGGCAGTATTTGTGG |
| HAPI1_4_API5_F_AG | aatactacctgctGAATGTGCTGTTTTTGAGAC |
| hapi1MID_apiWH2_R_AG | ccatgagcgaCTCAGGTGAGACTAAATGATC |
| apiMID_hapiWH2_F_AG | attgggtattTCACCACCAAGTACAGAAATTG |
| hapi1WH2_pl_R_AG | tcgagggtacctctagacccTCAAGAATCACTCCAACTG |
| SB295 | GGGGACAAGTTTGTACaaaaaagcaggctATGTCAGATGTAAATAGTTCTTCCAAG |
| SB296 | GGGGACCACTTTGTAcaagaaagctgggttTCACTTGTACAGCTCGTCCATG |
| SB323 | GGGGACAAGTTTGTACaaaaaagcaggctATGGGAGGTTCTGGTGGAGGTGGATCAG |

| | |
|---|---|
| SB329 | GGGGACAAGTTTGTACaaaaaagcaggctATGTCAGATGTAAGTTCTTCCAAGAAAGATAAATC |
| SB324 | GGGGACAAGTTTGTACaaaaaagcaggctATGAGCACTGTTGTTGAAAATACACAATC |
| SB325 | GGGGACAAGTTTGTACaaaaaagcaggctATGGAACCCATTTTATTCACAACTTC |
| SB343 | GGGGACAAGTTTGTACaaaaaagcaggctatgAGCACTGTTGTTGAAAATACACAATCAGAAATTGAAGGTACACCATCGAACC |
| SB342 | GGGGACAAGTTTGTACaaaaaagcaggctATGTCAGATGTAAATAGTTCTTCCAG |
| SB268 | AAGCTGACCCTGAAGTTCATCTGC |
| SB269 | CTTGTAGTTGCCGTCGTCCTTGAA |
| SB304 | AAGGATGCCGTGAAGAAGATGT |
| SB305 | GCATCGTAGTCAGGAGTCAACC |
| SB306 | GGCACTCACAAACGTCTATTTC |
| SB307 | ACCTGGGAGGCATCCTGCTTAT |

**Table S1: Primer sequences mentioned in this study**

**Table S2**

| Name | Source | Description |
|---|---|---|
| pKGW_RR_MGW | Gavrin et al., 2020 | Whole vector sequence of multisite Gateway-compatible destination vector with DsRed casette |
| pENTR4_1_prAtUBQ3p | Gavrin et al., 2020 | Whole vector sequence of pENTR vector with Arabidopsis UBQ3 (Gavrin ref) flanked by attL4 and attR1 sites |
| pENTR4_1_prMtAPI | Gavrin et al., 2020 | Whole vector sequence of pENTR vector with 2kb Medicago API promoter (Gavrin ref) flanked by attL4 and attR1 sites |
| pENTR_p2rp3_T35STerm | Gavrin et al., 2020 | Whole vector sequence of pENTR vector with T35S terminator flanked by attR2 and attL3 sites |
| API | Gavrin et al., 2020 | Coding sequence of Medicago API (Medtr4g013235) |
| HAPI1 | Gavrin et al., 2020 | Coding sequence of Medicago HAPI1 Medtr7g071440 |
| API_H2 | This paper | Coding sequence of API chimera |
| API_H4 | This paper | Coding sequence of API chimera |
| API_H2+4 | This paper | Coding sequence of API chimera |
| HAPI1_A2 | This paper | Coding sequence of HAPI1 chimera |
| HAPI1_A4 | This paper | Coding sequence of HAPI1 chimera |
| HAPI1_A2+4 | This paper | Coding sequence of HAPI1 chimera |
| API_mCitrine | This paper | Coding sequence of API fusion |
| API_delABC_mCitrine | This paper | Coding sequence of API variant fusion |
| API_GS_mCitrine | This paper | Coding sequence of API variant fusion |
| API_delA_mCitrine | This paper | Coding sequence of API variant fusion |
| API_delB_mCitrine | This paper | Coding sequence of API variant fusion |
| API_delC_mCitrine | This paper | Coding sequence of API variant fusion |
| API_delAB_mCitrine | This paper | Coding sequence of API variant fusion |
| API_delBC_mCitrine | This paper | Coding sequence of API variant fusion |
| API_K517R_mCitrine | This paper | Coding sequence of API variant fusion |
| API_K518R_mCitrine | This paper | Coding sequence of API variant fusion |
| API_K520R_mCitrine | This paper | Coding sequence of API variant fusion |
| API_K547R_mCitrine | This paper | Coding sequence of API variant fusion |
| API_3xKtoR_mCitrine | This paper | Coding sequence of API variant fusion |
| API_K4xKtoR_mCitrine | This paper | Coding sequence of API variant fusion |
| HAPI1_mCitrine | This paper | Coding sequence of HAPI1 fusion |
| HAPI1_insABC_mCitrine | This paper | Coding sequence of HAPI1 variant fusion |
| HAPI1_GS_mCitrine | This paper | Coding sequence of HAPI1 variant fusion |
| 42aa_mCitrine | This paper | Coding sequence of mCitrine variant fusion |
| GS_mCitrine | This paper | Coding sequence of mCitrine variant fusion |

| | | |
|---|---|---|
| 41aa_mCitrine | This paper | Coding sequence of mCitrine variant fusion |
| 2-20aa_mCitrine | This paper | Coding sequence of mCitrine variant fusion |
| 22_43aa_mCitrine | This paper | Coding sequence of mCitrine variant fusion |
| 31_43aa_mCitrine | This paper | Coding sequence of mCitrine variant fusion |
| 22_30aa-mCitrine | This paper | Coding sequence of mCitrine variant fusion |
| 4xSTtoA_mCitrine | This paper | Coding sequence of mCitrine variant fusion |
| 4xSTtoD_mCitrine | This paper | Coding sequence of mCitrine variant fusion |
| 7xSTtoA_mCitrine | This paper | Coding sequence of mCitrine variant fusion |
| 7xSTtoD_mCitrine | This paper | Coding sequence of mCitrine variant fusion |
| 9xSTYtoA_mCitrine | This paper | Coding sequence of mCitrine variant fusion |
| 9xSTYtoD_mCitrine | This paper | Coding sequence of mCitrine variant fusion |
| 4xKtoR_mCitrine | This paper | Coding sequence of mCitrine variant fusion |

**Table S2: Description of vector and coding sequences mentioned in this study.**

**Data S1. (separate file)**

**Data S1: Fasta file of vector and coding sequences mentioned in this study**
